# Gut microbiota influence B cell function in a TLR5-dependent manner

**DOI:** 10.1101/537894

**Authors:** Sha Li, William A. Walters, Benoit Chassaing, Benyue Zhang, Qiaojuan Shi, Jillian Waters, Andrew T. Gewirtz, Cynthia A. Leifer, Ruth E. Ley

## Abstract

Toll-like receptor (TLR) 5-deficient mice display aberrantly low levels of flagellin-specific antibodies (Flic-IgA) secreted into the gut, combined with excess bacterial flagellin in the gut, and together these attributes define microbiome dysbiosis (T5-dysbiosis). How TLR5 signaling deficiency results in T5-dysbiosis is unclear. Here, we address the role of B cells in T-dysbiosis. We observed that B cells do not express TLR5, and that B cell transplantation from TLR5^-/-^ mouse donors into B-cell deficient mice resulted in a slight reduction in Flic-IgA levels compared to B-cells from WT donors. Bone marrow transplants from WT and TLR5^-/-^ donors into recipients of both genotypes confirmed that TLR5 signaling by non-hematopoietic cells is required for T5-dysbiosis. We observed TLR5 deficiency was associated with an expanded population of IgA+ B cells. TLR5^-/-^ mice tended to have higher richness for the IgA gene hypervariable region (CDR3 gene) variants. Transplantation of microbiomes from TLR5^-/-^ and WT microbiomes donors into germfree mice resulted in a higher proportion of IgA-secreting B cells, and higher overall fecal IgA and anti-Flic IgA for TLR5^-/-^ microbiome recipients. This observation indicated that the TLR5^-/-^ mouse microbiome elicits an anti-flagellin antibody response that requires TLR5 signaling. Together these results indicate that TLR5 signaling on epithelial cells influences B cell populations and antibody repertoire.

## Introduction

The bacterial flagellum is comprised of polymerized flagellin, a highly conserved monomeric protein that is specifically recognized by the mammalian Toll-like receptor (TLR) 5 (Ramos et al., 2004; Smith et al., 2003). TLR5 is highly expressed basolaterally on intestinal epithelial cells, as well as by dendritic cells in the gut (Gewirtz et al., 2001; Leifer et al., 2014; Lim et al., 2015; Uematsu et al., 2006). TLR5 has an important role in the inflammation response to flagellated pathogens that breach the epithelial barrier. By triggering TLR5, flagellin initiates the NF-κB-mediated production of inflammatory cytokines and chemokines (Didierlaurent et al., 2004; Eaves-Pyles et al., 2001; Hayashi et al., 2001; Tallant et al., 2004; Yu et al., 2004). In addition to its role in defense against pathogens, TLR5 signaling also participates in the pathology of Crohn’s disease (Gewirtz et al., 2006; Lodes et al., 2004).

Loss of TLR5 signaling impacts the commensal microbiota and alters intestinal homeostasis ((Carvalho et al., 2012; Singh et al., 2015; Vijay-Kumar et al., 2010)). TLR5-deficient (TLR5^-/-^) mice exhibit metabolic abnormalities, and are predisposed to colitis: both syndromes are dependent on elevated levels of inflammatory microbe-associated molecular patterns (MAMPs) in the TLR5^-/-^ gut microbiome (Carvalho et al., 2012; Singh et al., 2015; Vijay-Kumar et al., 2010). TLR5^-/-^ mice harbor gut microbiota with higher biomass levels and higher levels of bacterial bioactive flagellin compared to WT mice, regardless of the diversity of the microbiota or the provenance of the mice (Cullender et al., 2013; Vijay-Kumar et al., 2010). Furthermore, a higher bioactive flagellin load in the TLR5^-/-^ mouse gut is associated with lower levels of flagellin-specific IgA despite higher overall levels of circulating and intestinal IgA, compared to WT (Carvalho et al., 2012; Cullender et al., 2013; Singh et al., 2015; Vijay-Kumar et al., 2010). Thus, TLR5-deficient mice display host-microbial dysbiosis (hereafter, T5-dysbiosis) characterized by abnormally high levels of total IgA and flagellin in the gut, combined with abnormally low anti-flagellin IgA.

Expression of flagellin by intestinal bacteria is induced according to local conditions encountered in the gut, and there is some evidence that the presence of antibody in the gut can affect flagellin expression. For instance, we previously showed that a lack of secreted IgA (sIgA) in the mouse gut (i.e., *Rag1*^-/-^ mice) was associated with elevated levels of flagellin production by *E.coli* in a tri-colonized gnotobiotic model (Cullender et al., 2013). In vitro, direct administration of anti-Flic IgA to *E.coli* can reduce flagellin gene expression (Cullender et al., 2013). Together, these observations support the hypothesis that anti-Flic IgA in the gut can serve to regulate flagellin expression. The mechanisms underlying the shifts in the production of total IgA and of flagellin-specific in TLR5-deficient hosts are not well understood.

Loss of TLR5 signaling on intestinal epithelial cells is sufficient to partially recapitulate the effects of systemic TLR5-deficiency (Chassaing et al., 2014). Nevertheless, B cells transferred from MyD88-deficient mice (e.g., with attenuated signaling from TLR5) into MT mice (that lack endogenous B cells) has resulted in phenotypes characteristic of the T5-dysbiosis, e.g., reduced anti-Flic IgA in serum (He et al., 2007; Oh et al., 2014; Pasare and Medzhitov, 2005). These findings imply that reduction in the level of Flic-specific IgA in TLR5^-/-^ mice could result from the loss of TLR5 signaling directly on B cell function, or be a consequence of compromised B cell development in a TLR5-deficient host. Here, we explored the role of B cells on the TLR5^-/-^ associated T5-dysbiosis.

## Results

We first examined the metabolic phenotype and gut homeostasis for our in-house TLR5^-/-^ mouse colony. To reduce the bias caused by co-caging and littermate effects, which are know to impact the microbiota (Goodrich et al., 2014), we randomly selected age-matched WT and TLR5^-/-^ male littermates derived from different sets of breeders that were not co-housed after weaning and examined their phenotypes (methods). Compared to WT littermates, TLR5^-/-^ mice displayed slightly but significantly greater body weight (Fig 1a). Although the difference in body mass between the genotypes was small but significant, the epididymal and mesenteric fat masses nearly double for the TLR5^-/-^ mice (Fig 1b-c). TLR5^-/-^ mice also displayed reduced glucose tolerance (Fig 1d). Stool from TLR5^-/-^ mice contained higher levels of total IgA and bioactive bacterial flagellin (Fig 1e,f), and reduced levels of IgA antibody reactive with recombinant *E. coli* Flic protein (Fig 1g), compared to WT mice. We obtained similar patterns when examining these parameters on age-matched WT and TLR5^-/-^ male non-littermates. In all, these data demonstrated that mice in our TLR5^-/-^ colony recapitulated previously reported metabolic and T5-dysbiosis phenotypes described for mice with different founders (Cullender et al., 2013; Vijay-Kumar et al., 2010).

**Fig. 1:**
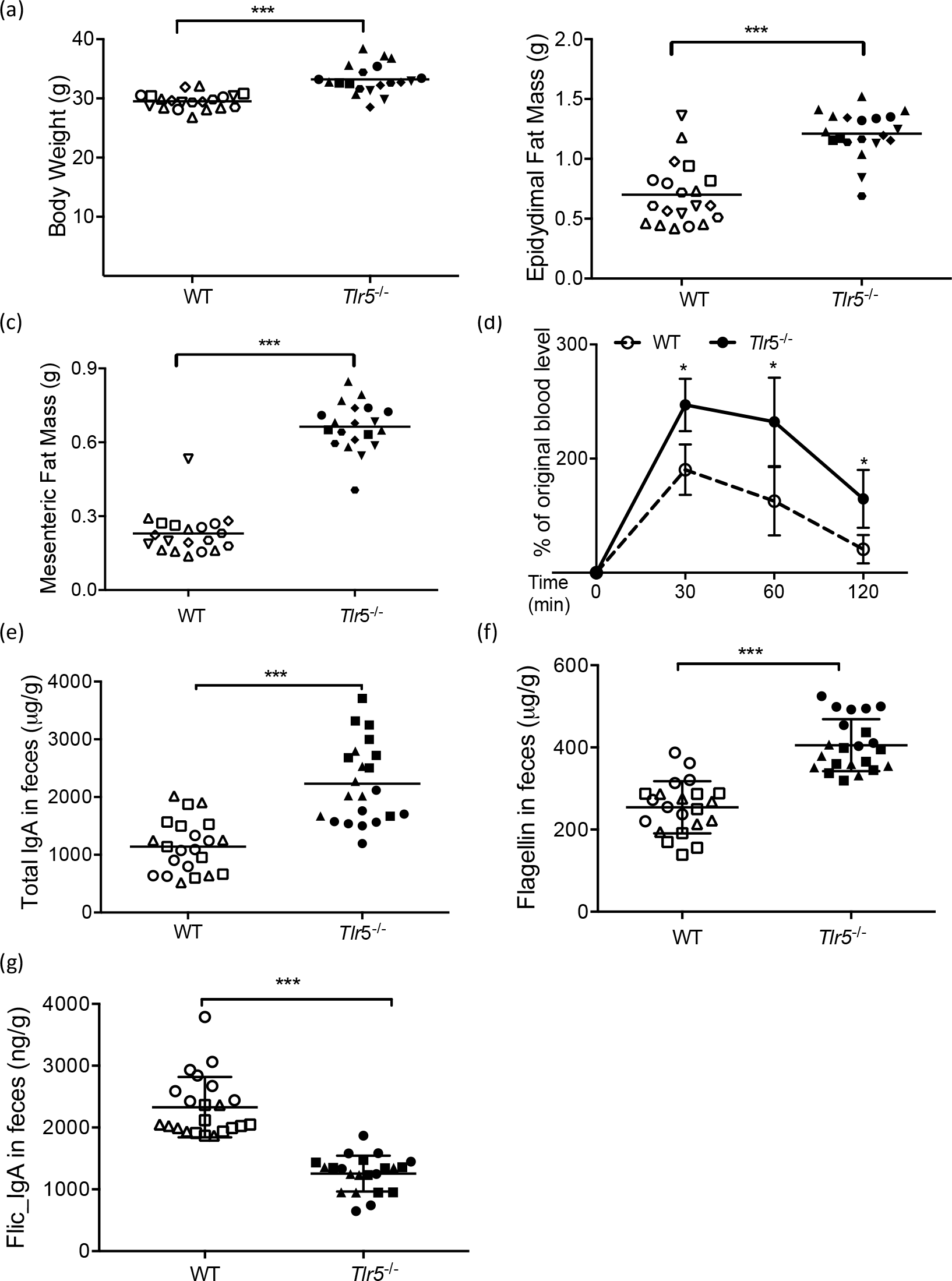
TLR5^-/-^ mice display metabolic syndrome and altered levels of antibodies and flagellin. (a) Body weight. (b) Epididymal fat pad mass. c) Mesenteric fat mass. The data were pooled from six independent assays with totally n = 20 for each group (for a-c). (d) Glucose tolerance test. Data were pooled from three independent assays with n = 9 for each group. Age-matched WT and TLR5^-/-^ male littermates were used in these assays. (e) Total fecal IgA. (f) Flagellin in feces. (g) Fecal Flic-specific IgA. Data from the same assay are depicted with the same symbol. Data from age-matched WT and TLR5^-/-^ male littermates are represented as circle, square, diamond, and hexagon; while data from age-matched WT and TLR5^-/-^ male non-littermates are depicted as triangles and inverted triangles. Data were pooled from three independent assays with n = 22 for each group. *: p<0.05; ***: p<0.0001.

### B cells do not express TLR5

Whether or not TLR5 is expressed on B cells remains controversial (Genestier et al., 2007; Gururajan et al., 2007; He et al., 2007; Pasare and Medzhitov, 2005). We first examined the expression of TLR5 on B cells by flow cytometry. In accord with recent studies, we detected TLR5 expression on neutrophils from WT but not TLR5^-/-^ mice, demonstrating the specificity of the anti-TLR5 antibody (Figure 2a). In contrast, this antibody failed to stain B cells isolated from WT bone marrow, spleen, ileum, or colon (Figure 2b-c). Thus, we conclude that mouse B cells do not express TLR5, excluding the possibility that cis-regulated TLR5 signaling in B cells is responsible for T5-dysbiosis.

**Fig. 2:**
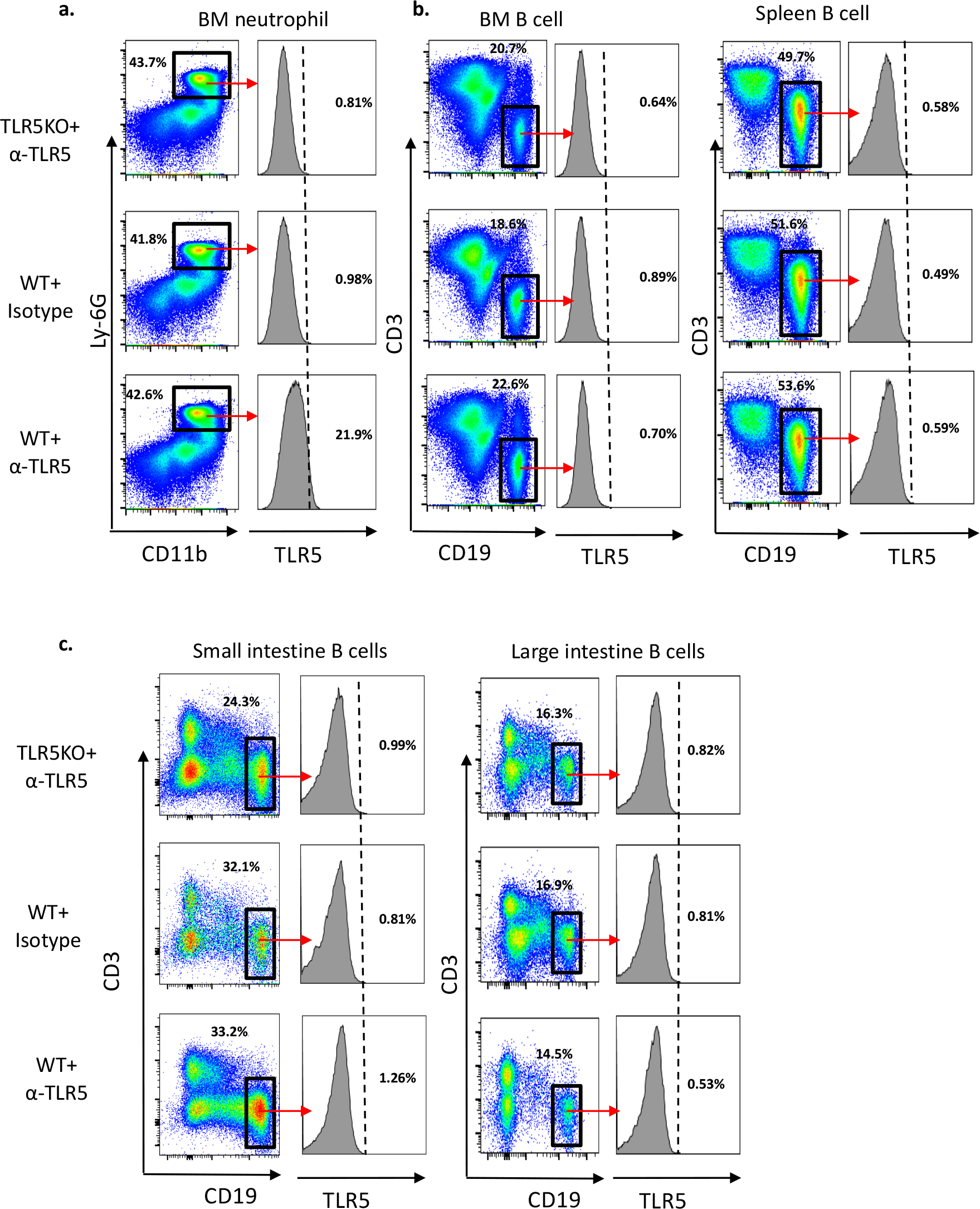
No detectable TLR5 on B cells. Cells were isolated from different tissues of WT and TLR5^-/-^ mice. TLR5 expression on neutrophils (CD11b^+^Ly6G^+^) (a) and B cells (CD3^−^CD19^+^) from various organs (b, c) depicted by histogram plots. BM = bone marrow. Mouse IgG2a was used as isotype control. These are representative results of three experiments.

### The B cell developmental environment can induce a partial T5-dysbiotic phenotype

To test whether T5-dysbiosis could be the consequence of TLR5 deficiency impaired B cell development, we purified WT and TLR5^-/-^ B cells from age-matched male littermates and transferred them into B cell-deficient MT male mice (Fig 3a). The recipient mice were single-housed ten days prior to B cell transfer and maintained in the same cage during the whole experimental period. We observed that 12 days post-transfer, the transfer of B cells derived from WT donors into MT mice had resulted in significantly higher levels FliC-IgA in feces compared to B cells derived from TLR5^-/-^ donors (Fig 3b). There were no differences in body weight, total fecal IgA, or flagellin levels between groups at either 7 or 12 days post-transfer (Fig 3c-e). Thus, the B cells that developed in a TLR5^-/-^ host environment had a reduced capacity to produce antibody against flagellin compared to WT, suggesting that the WT environment induced a larger clonal expansion of B cells producing anti-flagellin antibody. However, the reduction of anti-flagellin IgA in recipients of B cells from TLR5^-/-^ hosts compared to WT hosts was not sufficient to alter flagellin levels in the gut, within the 12-day period.

**Fig. 3:**
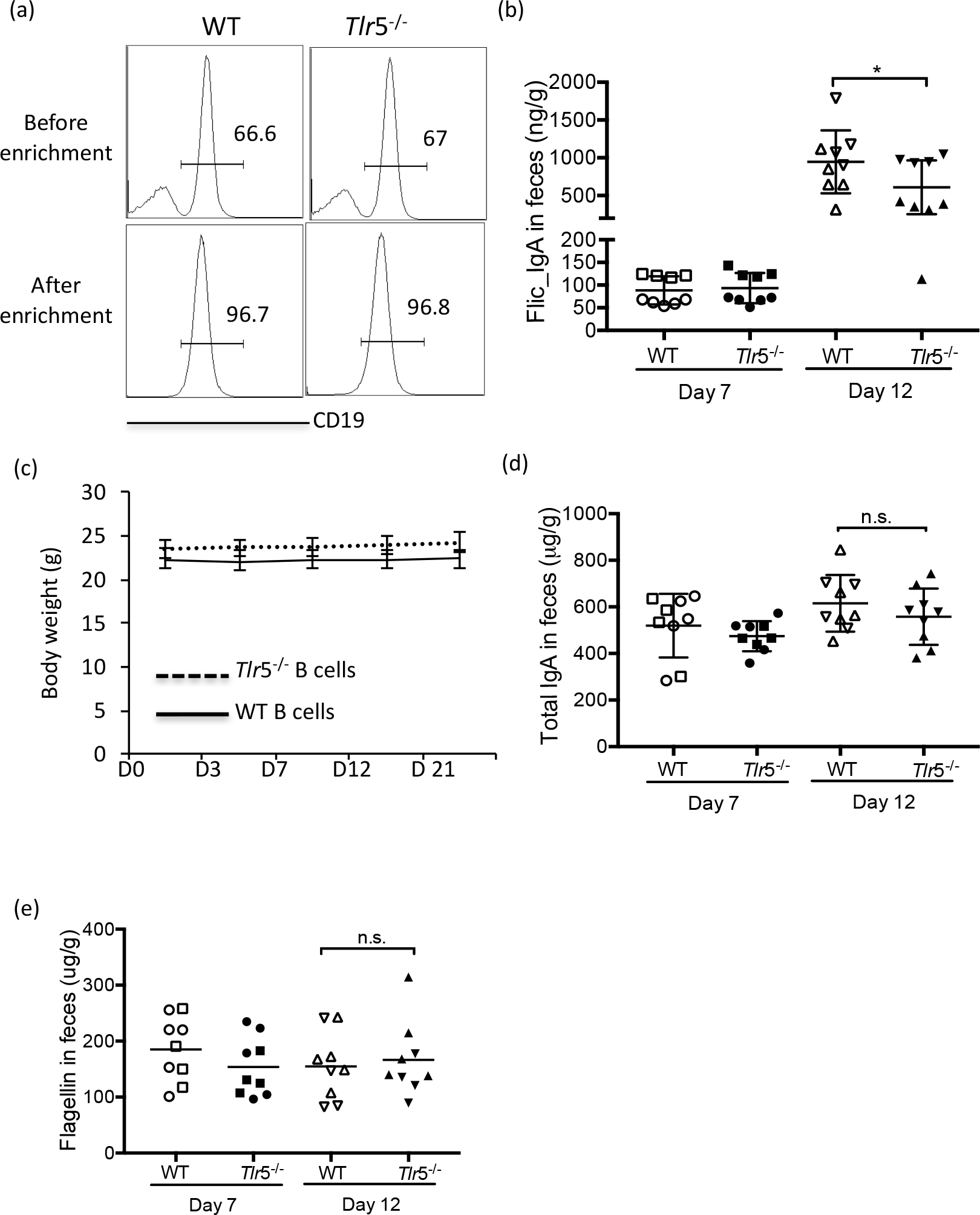
Transfer of splenic TLR5^-/-^ B cells did not result in TLR5 deficiency-induced phenotype. Splenocytes were isolated from 12-week old WT and TLR5^-/-^ male littermates. B cells were enriched and transferred into seven-week-old male μMT mice. (a) The percentages of isolated splenic B cells before (top) and after (bottom) enrichment. Data were representative of two independent assays. Flagellin specific IgA (b), body mass (c), total IgA (d), and flagellin levels (e) were measured. Data from the same assay were represented with the same symbol. Dotted line: μMT mice receiving splenic *Tlr5*^-/-^ B cells; solid line: μMT mice receiving splenic WT B cells. Data are pooled from two independent assays with total n = 9 for each group. *: p<0.05; n.s.: not significant.

### TLR5 deficiency on hematopoietic cells is insufficient to induce T5-dysbiosis and has a minimal effect on the microbiota

The cell types known to express TLR5 include epithelial cells, dendritic cells and T cells (Crellin et al., 2005; Descamps et al., 2012; Gewirtz et al., 2001; Lim et al., 2015; Oh et al., 2014; Pasare and Medzhitov, 2005; Sandig and Bulfone-Paus, 2012; Uematsu et al., 2006). To further investigate the cellular targets of TLR5 signaling during homeostasis, we established chimeric mice by reconstituting sublethally irradiated non-littermate WT and TLR5^-/-^ recipients with either WT or TLR5^-/-^ donor bone marrow cells (Fig 4a). We used age-matched WT and TLR5^-/-^ males as donors and recipients. Recipient mice of the same genotype were derived from different breeders, received the same source of bone marrow cells, and were housed together during the experimental period (methods). We performed two independent replicates of this experiment (replicate results shown in Supplemental Fig 1). We monitored host weight and collected fecal samples twice per week. Note that in the TLR5^-/-^ donor to WT recipient treatment, in both replicates of the experiment, a slight contamination of WT bone marrow was evident from a genotype-specific PCR assay performed on DNA purified from blood at week 4 post-transfer (Supplemental Fig 2).

**Fig. 4:**
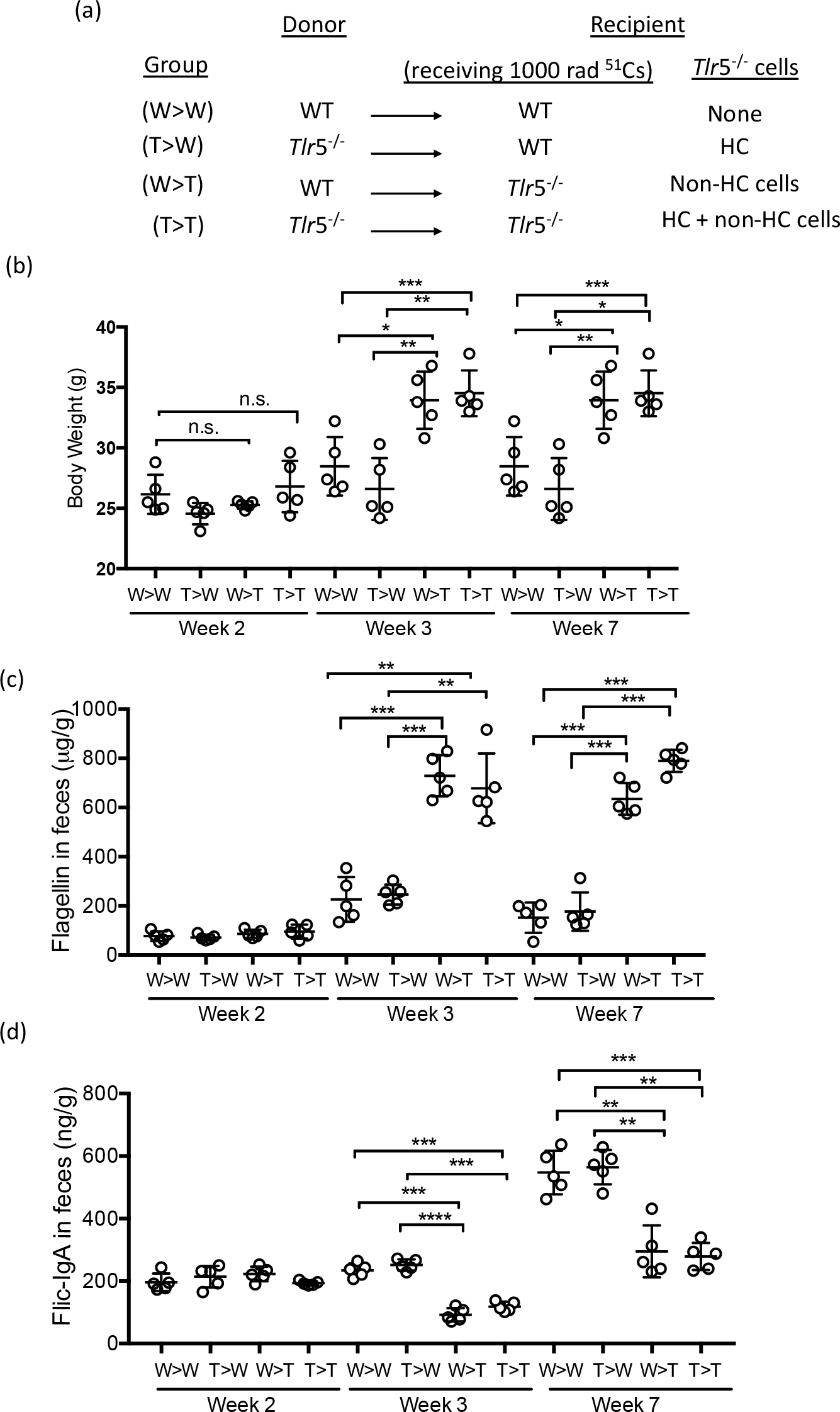
TLR5 on non-hematopoietic cells is responsible for T5-dysbiosis. (a) Schematic of the bone-marrow transfer assay. Age-matched WT and *Tlr5*^-/-^ non-littermates were used for both donors and recipients in this experiment. N= 5 for each chimera group. HC=hematopoietic cells. (b) Body weight. (c) Fecal flagellin level. (d) Fecal Flic-specific IgA concentration. *: p<0.05; **: p<0.01; ***: p<0.0001; n.s.: not significant. Repeated experiment results shown in Supplemental figure 3.

We observed that TLR5^-/-^ mice that received bone marrow from either donor (e.g., WT >TLR5^-/-^ and TLR5^-/-^ >TLR5^-/-^), displayed a significant increase in body weight, compared with WT recipients that received bone marrow from either donor (e.g., WT>WT and TLR5^-/-^ >WT; Fig 4b) 3 and 7 weeks post-transfer, that was not evident 2 weeks post-transfer. Levels of bioactive flagellin in feces were comparably low in WT recipients of both donor types, and higher in all TLR5^-/-^ recipients at three and seven weeks post-transfer (Fig 4c). Note that the low flagellin in both WT and TLR5^-/-^ recipients at 2-weeks post-transfer may be the results of the antibiotic treatment (for one week post-irradiation, see Methods).

Flic-IgA levels in feces were similar across treatments at week 2 (Fig. 4d). At week 3 post-transfer, the TLR5^-/-^ recipients saw a drop in anti-Flic IgA that was not present in the WT recipients. At week seven, WT recipients showed a significant increase in Flic-IgA levels in feces, whereas levels in the TLR5^-/-^ recipients returned to the 2-week baseline (Fig. 4d). These results support a role of non-hematopoietic cell-specific TLR5 in maintaining normal levels of anti-flagellin IgA secreted to the gut.

### Effects of bone marrow transplant on microbiota

To evaluate the effects of the bone marrow transfer treatments on the gut microbiota, we characterized the fecal microbial diversity of the bone marrow recipients by sequencing the V4 region of 16S rRNA gene PCR-amplicons derived from bulk fecal DNA samples at different time points. Data from the two independent replicates were pooled; 16S rRNA gene sequences clustered them into 97% ID OTUs (operational taxonomic units). After quality filtering and discarding of singletons, we obtained 78,502 (mean) ± 19,543 (standard deviation, S.D.) sequences per sample. We first analyzed richness (alpha diversity) in fecal samples from all time points after bone marrow transfer (see Methods). Alpha-diversity results were similar, with no significant differences for all samples based on phylogenetic diversity and observed OTUs calculations, indicating that neither donor bone marrow sources or recipient genotypes affected fecal microbiota richness (Supplemental Figure 3a).

A common approach to microbiome analysis is to generate distance metrics between samples and search for patterns related to experimental treatments. In this case, the co-caging of mice was confounded with experimental treatments. Samples, with unweighted or weighted UniFrac distances, clustered according to donor and recipient genotype status, which also matches co-caging status, and no age gradient is apparent (Figure 5, Supplemental figure 3 b-d). Therefore, we used a linear mixed model approach (Methods) to identify OTUs whose relative abundances differed between treatments (Supplemental table 1). Results showed that several OTUs belonging to the Clostridiaceae family or the Bacteroidales (S24-7 candidate family) were significantly enriched in the WT recipient mice. A Mollicutes (RF39 candidate order) OTU was enriched in the Tlr5^-/-^ recipient mice. It is not possible to tell from these taxonomic annotations if the bacteria enriched in either host genotype are flagellated. Overall, the result of these analyses indicate that the composition of the fecal microbiome post-bone marrow transfer was most greatly influenced by the recipient’s genotype.

**Fig. 5:**
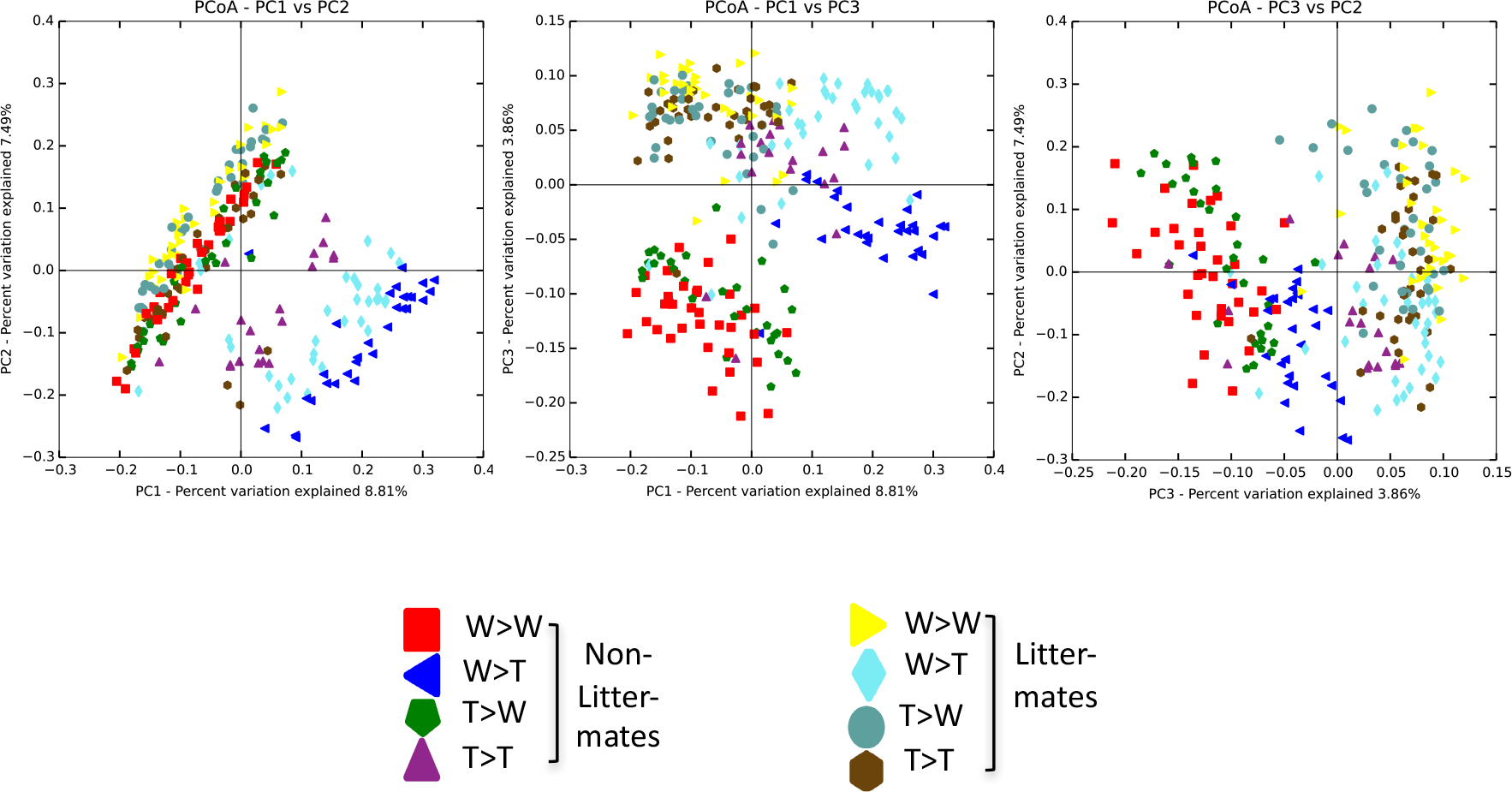
Chimeric mouse gut metagenomes cluster according to donor and recipient genotype status, which also matches co-caging status. Clustering of samples with unweighted UniFrac distances. The first three principal coordinates are plotted, with percent variance explained displayed on the axes. Samples are labeled to indicate donor and recipient genotypes (W = wild type, T = TLR5^-/-^). Color and shape indicate litter-mate status.

### TLR5 deficiency leads to the expansion of IgA-secreting B cells and upregulation of B cell activating factor

An elevated level of total IgA in TLR5^-/-^ fecal samples indicates a potential increase in IgA-secreting B cells. We examined intracellular IgA abundance in mononuclear cells isolated from the *lamina propria* of the large intestines of WT and TLR5^-/-^ mice (Figure 6, Supplemental Fig 4). To eliminate any confounding effects of husbandry, we housed littermates from multiple sets of breeders separately based on the genotype and their progenitors. The total cell numbers isolated from the *lamina propria* of the large intestine were comparable between WT and TLR5^-/-^ mice (n=8/group; WT: 1.76 ± 0.40 × 10^6^ and *Tlr5*^-/-^: 1.84 ± 0.59 × 10^6^). In contrast to the similar proportion of splenic IgA-secreting cells between WT and TLR5^-/-^ mice, we detected a significantly higher proportion of IgA^+^ B cells in the lamina propria of TLR5^-/-^ large intestine (Fig 6a, b).

**Fig. 6:**
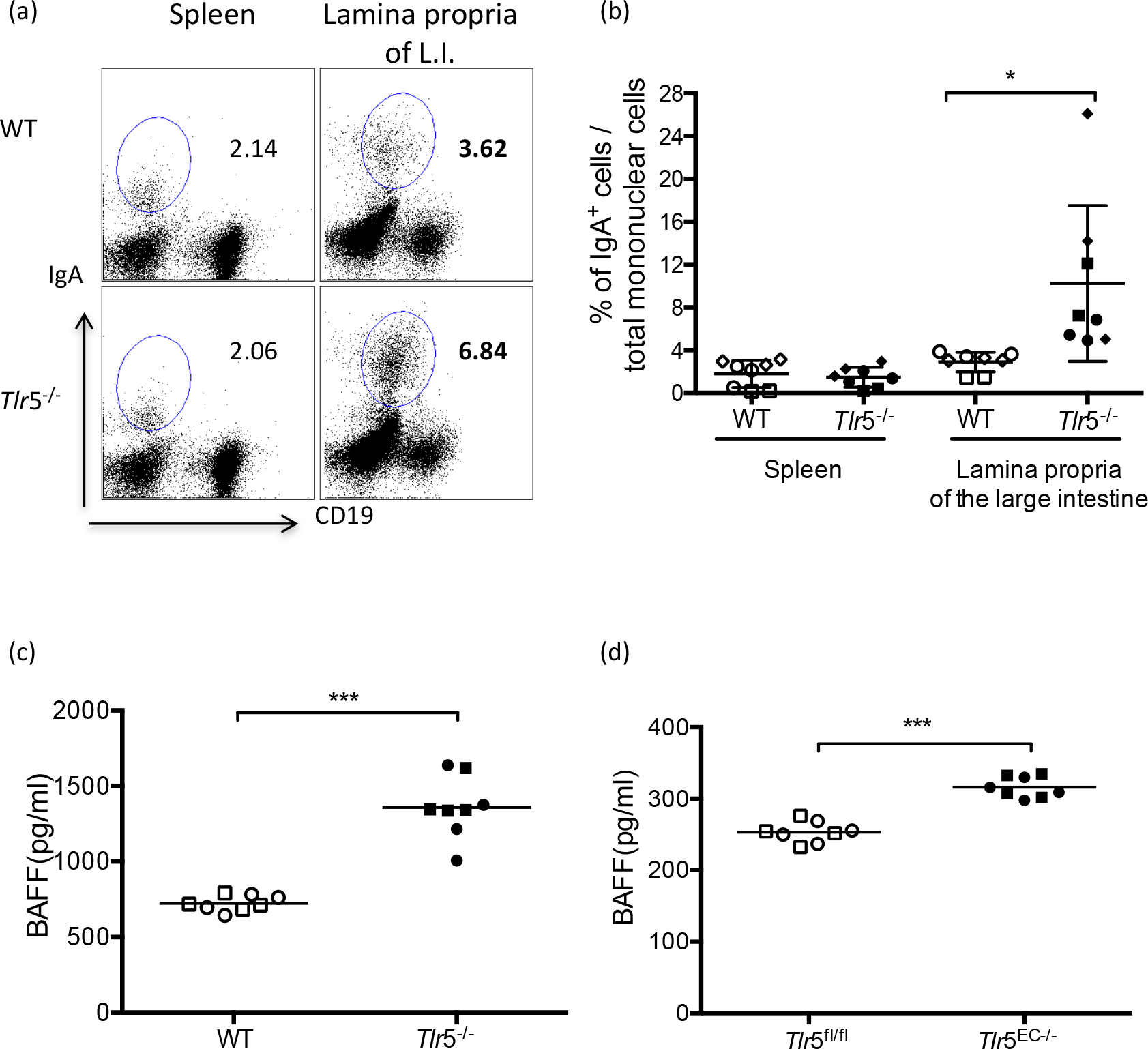
TLR5 deficiency enriches IgA-secreting B-cells and augments BAFF levels. (a-b) Percentages of IgA^+^ cells in the spleen and lamina propria of the large intestine. Cells were isolated and stained with the B cell surface marker, CD19, and intracellular IgA mAb. (a) Representative data of the percentages of IgA^+^ cells/total mononuclear cells. (b) The percentages of IgA^+^ cells were pooled from three independent assays and plotted with n= 8 for each group. (c-d) BAFF levels in the homogenate supernatant of the large intestines from age-matched WT and *Tlr5*^-/-^ male littermates (c), or *Tlr5*^fl/fl^ and *Tlr5*^EC−/−^ male littermates (d). Data were pooled from two independent assays with n = 8 for each group. In (b-d), each symbol represents an individual sample. Data from the same assay were depicted with the same shape of the symbol. *: p<0.05; ***: p<0.0001

BAFF (B-cell activating factor) is widely known to regulate B cell proliferation and class switch recombination to IgA (Litinskiy et al., 2002). To investigate the relationship of BAFF to the large intestinal IgA^+^ B cells, we examined BAFF levels in the large intestine of age-matched WT and *Tlr5*^-/-^ male littermates. We observed a higher level of BAFF in TLR5^-/-^ large intestines compared WT samples (Fig 6c). Furthermore, as intestinal epithelial-specific TLR5-knockout (*Tlr5*^EC−/−^) mice have been shown to exhibit many features of the T5-dysbiosis (Chassaing et al., 2014), we compared BAFF levels in the large intestines of TLR5^EC−/−^ mice with those of TLR5^fl/fl^ siblings expressing flanking loxP, but not the Cre gene. We observed a higher level of BAFF in the large intestine of TLR5^EC−/−^ mice, further indicating the trans-effect of microbiome-induced intestinal epithelial cell-specific BAFF on intestinal B cell function (Fig 6d). Thus, BAFF may be one of the factors leading to elevated proportion of IgA+ B cells in the TLR5^-/-^ large intestine *lamina propria*. (Note that the BAFF measures should be normalized between genotypes (by intestine weight, or number of Peyer’s patches) for a more robust comparison.)

### Transplant of WT and TLR5^-/-^ mouse microbiomes into germfree recipients leads to differential B cell responses

Fecal transplants of the TLR5^-/-^ mouse microbiome into germfree WT recipients has previously been shown to transfer the T5-dysbiosis metabolic phenotype, but its impact on antibody levels and B cell populations has not been examined. The TLR5^-/-^ microbiome is characterized by a high flagellin load compared to the WT microbiome, and might be expected to generate a B cell response that differs from that of the WT microbiome. We transferred cecal microbiomes from 12-week old WT and TLR5^-/-^ male littermates into 3-4 week old germ-free Swiss Webster male littermates (Fig 7a). Donors and recipients were housed singly after weaning and fed a standard mouse chow *ad-libitum*. Three weeks after microbiota transfer, we observed a higher proportion of IgA-secreting B cells in the *lamina propria* of the large intestines of TLR5^-/-^ microbiota recipients, but not in the spleen, compared to those of WT microbiota-recipients (Fig 7b, c). In addition, the recipients of TLR5^-/-^ microbiome showed higher average levels of total IgA in feces compared to WT microbiome recipients (Fig 7d). A higher Flic-specific IgA was also observed in the recipients of TLR5^-/-^ microbiome (Fig 7e). Therefore, the TLR5^-/-^ microbiome, when transferred to the germfree WT host, stimulated a B cell response in the *lamina propria* characterized by an expansion of IgA+ B cells, and a greater total IgA load in the gut that included greater anti-flagellin IgA.

**Fig. 7:**
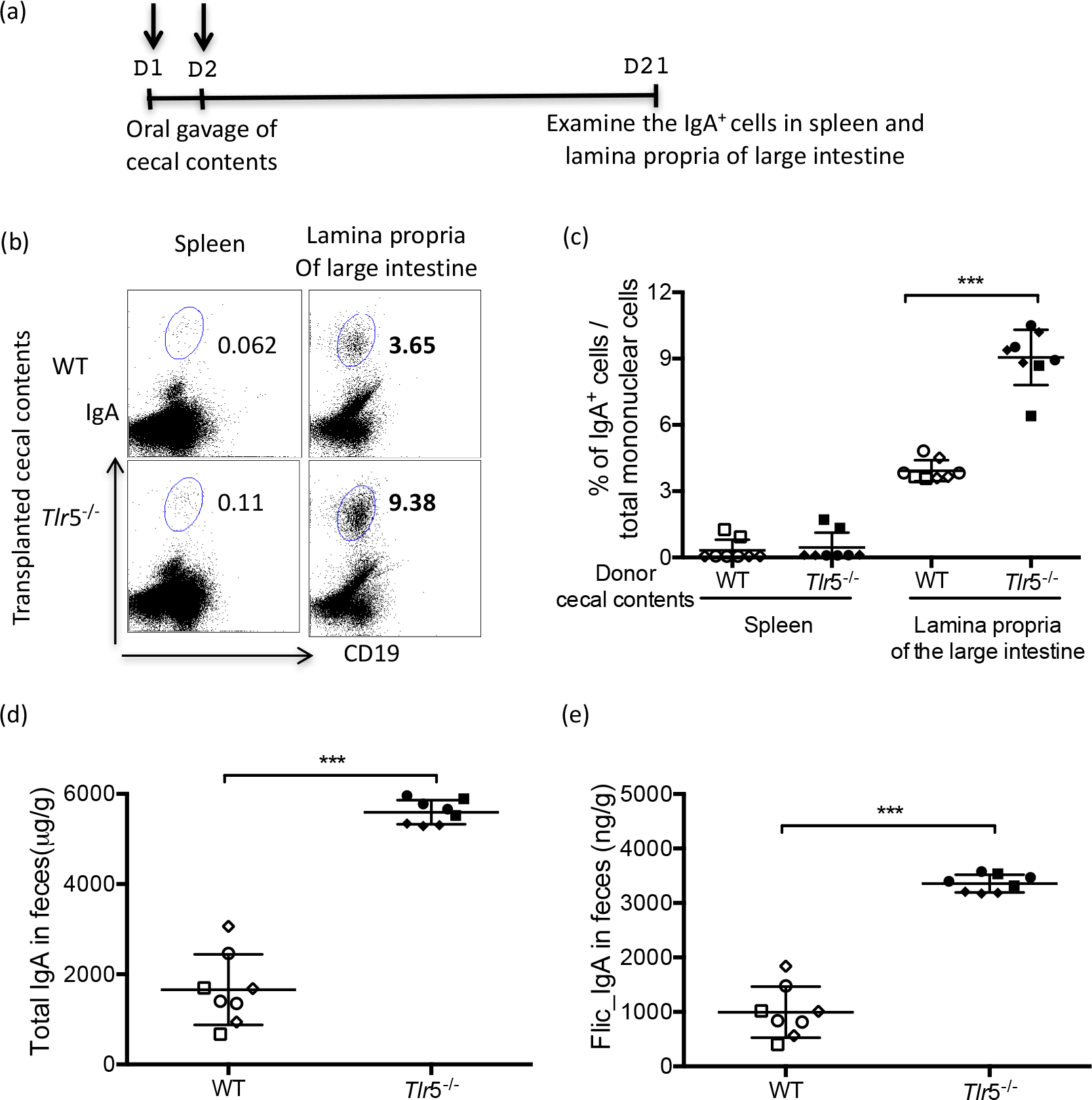
Transplant of WT AND TLR5^-/-^ cecal contents into germfree mice results in different IgA profiles. (a) Schematic of cecal transfer. Fresh cecal contents were collected from age-matched 10-12 week old WT and *Tlr5*^-/-^ male littermates and transferred into germ-free swiss-webster male littermates once daily for two consecutive days. Three weeks later, IgA^+^ cells were analyzed from the spleen and lamina propria. (b) Representative data of the percentages of IgA^+^ cells / total mononuclear cells. (c) Differences between percentage of IgA^+^ cells between recipient organ and donor genotype. Data were pooled from three independent assays with n = 8 for each group with data from the each assay were depicted with the same symbol.IgA (d) and Flic-specific IgA (e) were quantified from fecal samples, with each assay indicated by shared symbol. **: p<0.01; ***: p<0.0001

### TLR5-deficiency fine-tunes the IgA repertoire in the gut

To further compare intestinal B cell responses in TLR5^-/-^ and WT mice, we examined the IgA gene repertoire of the B cell populations derived from the spleen and the large intestine of WT (N=3) and TLR5^-/-^ (N=4) mice. We PCR-amplified the gene segments spanning the complementarity-determining region 3 (CDR3) of the IgA heavy chain, as this region is the most diverse and the key determinant of antigenic specificity (Davis et al., 1997; Xu and Davis, 2000) (PCR primers described in Supplemental Table 2). We obtained similar sequences counts for each sample (Supplemental Table 3).

We first examined richness (alpha-diversity) of CDR3 sequences based on the observed species (in this case, counts of unique IgA reads) and Shannon index (a measure of diversity and evenness) calculations. There was no difference in alpha-diversity between the spleen and large intestine IgA reads (Figure 8a). Within organs and between TLR5^-/-^ and WT genotypes, there was an observed increase in diversity in theTLR5^-/-^ samples, however these did have a significant p-value of less than 0.05.

**Fig. 8:**
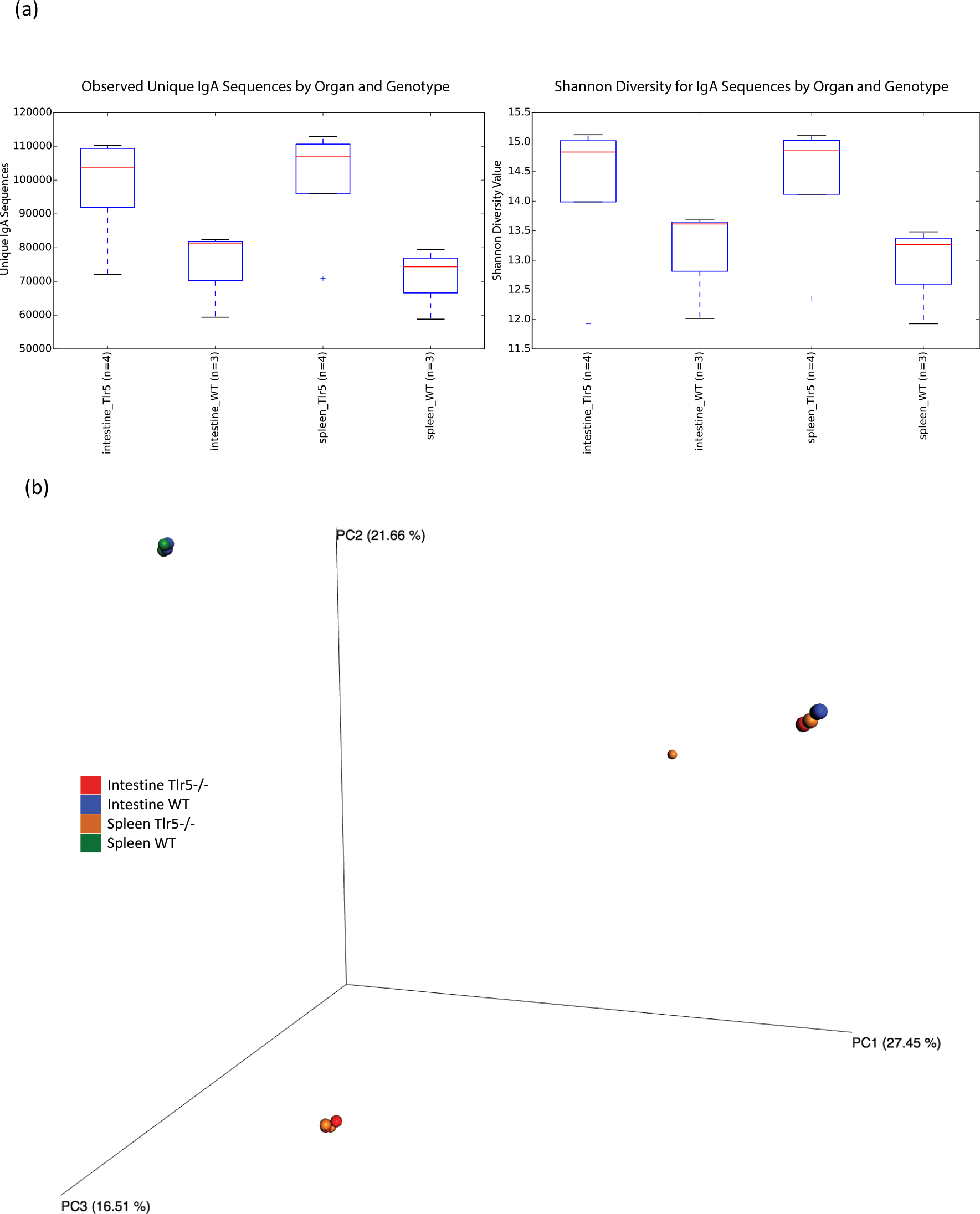
TLR5 deficiency increases overall diversity of IgA CDR3 sequences, while composition of IgA sequences is largely driven by individual mouse. Sequences of CDR3 regions of IgA heavy chain were compared between age-matched WT and TLR5^-/-^ male littermates (n=4 for TLR5^-/-^ organ/mice, n=3 for WT organ/mice).(a) Alpha diversity analysis. Sequences were clustered by the organs (spleen or the large intestine). Observed species (in this case, sequence variant; left) and Shannon diversity (right) were used to evaluate alpha diversity. (b) Beta diversity analysis. Bray-Curtis dissimilarity values were used to cluster samples, which show strong overlap as both the mice themselves in some cases and the between-organ IgA composition in all cases was very similar.

We also evaluated the dissimilarity of IgA CDR3 sequences from WT and TLR5^-/-^ mice based on the abundances of the unique sequences. We calculated Bray-Curtis distances between samples and applied principal coordinates (PC) analysis to the distance matrix. Plotting of the first three PCs showed that the samples from the same host clustered together, regardless of organ, indicating that a lot of similar sequences with high abundance were shared between organs within a single host (Figure 8b). To further corroborate this notion, we applied Adonis testing (Methods) to inspect the contributions of different variations (by mouseID, organ, mouse genotype, or the combination) to the sequence dissimilarities. Significant results were observed by mouse and genotype (Supplemental Table 4). Kruskal-Wallis tests for differential abundance of IgA sequences between organ and genotype showed no significant differences after multiple comparison corrections. Together these results indicate that the IgA CDR3 repertoires is not driven by organ type, but most strongly by the individual mice, with the genotype also exerting a secondary influence.

Unique IgA sequences were modeled into low and expanded frequency groups as described by Lindner (Lindner et al., 2012). The resulting rank versus frequency plots (Fig 9) showed a differential expansion between wild-type and TLR5^-/-^ mice, with a larger number of expanded frequency IgA clones in the TLR5^-/-^ mouse population (237 of 544 ranks were in the high frequency clone population for TLR5^-/-^ samples versus 179 of 589 ranks in the high frequency clone population for WT samples). Altogether, these data showed that TLR5 deficiency created a trend of higher diversity of CDR3 sequences and a larger pool of high-frequency IgA clones. These results may indicate that when TLR5 is present, flagellin becomes the dominant antigen thereby constraining B cell repertoire; when TLR5 is absent, there is less emphasis on flagellin, and thus more richness and diversity in the B cell repertoire.

**Fig 9:**
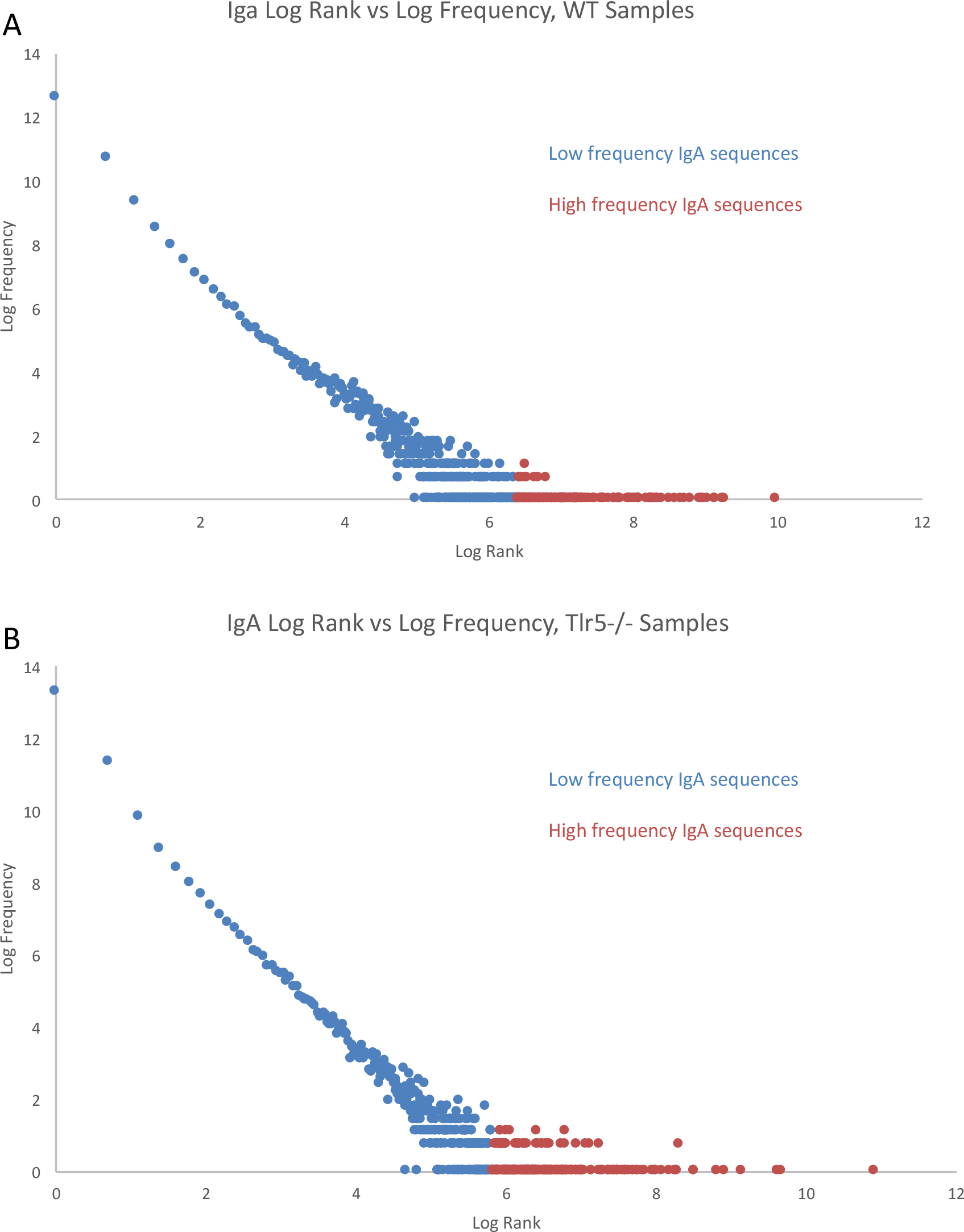
IgA CDR3 repertoire of WT and Tlr5^-/-^ mice. Log(rank) versus log(frequency) of wild type (a) and TLR5^-/-^ IgA CDR3 (B) sequences for spleen and large intestine samples combined. Low cell frequency populations are shown in blue and expanded cell populations are shown in orange.

## Discussion

TLR5 deficiency results in a T5-dysbiosis characterized by an elevated flagellin load in the gut and a skewed antibody profile, where greater than-normal levels of secreted IgA combine with lower-than-normal levels of anti-flagellin IgA. This T5-dysbiosis is also associated with metabolic inflammation. Given that the T5-dysbiosis is characterized in part by skewed antibody profiles, we asked about the role of B cells in driving the T5-dysbiosis. We found that B cells do not display TLR5 in either WT or TLR5^-/-^ hosts, and that IgA+ B cell populations are expanded in TLR5^-/-^ hosts and have an altered IgA gene repertoire. B cell and bone marrow transfers confirmed that TLR5 signaling on epithelial cells drives the T5 dysbiosis, possibly in part via elevated BAFF levels. Our results imply that B cells play a role in the T5-dysbiosis not due to intrinsic TLR5 signaling, but by conditioning through the responses of epithelial cells, that are themselves influenced by their own TLR5 signaling and by the responses of the microbiota.

TLR signaling may also impact B cell development by acting on their precursors in a trans-pattern: TLR signaling from other cell types may secrete mediators that regulate B cell development. In this situation, WT and TLR5^-/-^ B cells are likely to harbor distinct immune properties. To test this hypothesis, we transferred splenic WT and TLR5^-/-^ B cells into μMT mice that lack endogenous B cells. B cells conditioned in a TLR5+ environment resulted in higher Flic-IgA levels in recipient mice compared to B cell conditioned in the TLR5-host environment. This phenotype is consistent with the T5-dysbiosis, although we did not detect a difference in total IgA, which is typically associated with the T5-dysbiosis. Thus, the low Flic-IgA phenotype of the TLR5^-/-^ mouse could be transferred by the B cells derived from adult mice of that genotype.

We observed from our fecal transfers of TLR5^-/-^ and WT microbiomes into WT germfree mice that a microbiome with excess flagellin (TLR5^-/-^ donor) drives a higher Flic-IgA response. In the TLR5^-/-^ mouse, a high flagellin load is associated with a low FliC-IgA response despite high total IgA production, indicating a reduced capacity to produce this antibody. We checked the IgA gene (CDR3) repertoire for TLR5^-/-^ mice, which is directly linked to the diversity of IgAs, and can be influenced by the microbiota (Zeng et al., 2016). When comparing the CDR3 diversity of WT and TLR5^-/-^ mice, we observed a greater diversity for the TLR5^-/-^ mice. These data suggest that when TLR5 is present, flagellin is the dominant antigen.

The results of the bone marrow transfers performed here are consistent with a previous report that intestinal epithelial cell-specific TLR5 deficient mice partially recapitulate the phenotype of TLR5-null mice (Chassaing et al., 2014). Although the phenotype of chimera group TLR5^-/-^ to WT mimicked that of WT to WT chimera group, we detected PCR products of WT TLR5 gene in the blood lymphocytes in the group of TLR5^-/-^ to WT, such that some residual B cells may have been present. Nonetheless, the results from the three other chimeras strongly support the role of TLR5 signaling on epithelial cells in driving the T5-dysbiosis.

### Concluding remarks

Previous to this work, the role of B cells in the T5-dysbiosis was unclear. We showed here that TLR5-deficiency results in expansion of the B cell population, and that this expanded population carries a skewed IgA gene variant repertoire. Our work confirms the modest role of B cells in driving the T5-dysbiosis, and underscores the capacity of the microbiome to drive B cell responses.

## Methods

### Experimental Procedures

#### Mice

C57BL/6 (B6) WT mice, TLR5^-/-^ mice, and μMT mice were purchased from The Jackson Laboratory. Epithelial-specific TLR5^-/-^ mice were generously provided by Dr. Andrew Gewirtz’s lab (Georgia State University). These mouse strains were maintained and bred in Accepted Pathogen Barrier condition at Cornell University. 12-14 weeks old males were used in the experiments unless otherwise stated. Germ-Free (GF) Swiss Webster mice were purchased from Taconic Farms and bred in plastic gnotobiotic isolators at Cornell University. 3-4 weeks old GF male littermates were singly housed and used as recipients. All experiments involving mice were performed in accordance with the guidelines of the institutional animal care and use committee of Cornell University.

#### Breeding strategies

TLR5^-/-^ mice were crossed into the B6 background to produce F1 heterozygotes. F1 sister-brother matings were established to produce B6 and TLR5^-/-^ F2 homozygous littermates. After weaning at week 3, mice with the same genotype and progenitors were co-housed and used in all the experiments except bone marrow chimera assays. For mice used in bone marrow chimera assays, homozygous littermates of the same genotype, but from multiple breeders were housed together with 4-5 mice per cage after weaning. TLR5^-/-^ mice and WT mice were also bred separately to produce WT and TLR5^-/-^ non-littermates. All mouse colonies were maintained on breeder chow in the same room. The results obtained from littermates and non-littermates were depicted in different symbols in the figures.

#### Glucose tolerance test

Mice were fasted overnight for 15 hours. Baseline blood glucose levels were measured with an Accu-Check Advantage blood glucose meter (Roche) using tail vein blood. Glucose was injected into the intraperitoneal cavity at the concentration of 2g of glucose/kg body mass. The blood glucose levels were measured at 30, 60, and 120 minutes after injection.

#### Stool sample preparation

Fecal pellets were collected from individual mice and lyophilized (−20°C, 2 hours, atmospheric pressure 100Pa). Lyophilized products were suspended in PBS by homogenizing for 20 seconds with a Mini-Beadbeater-24 (BioSpec, Bartlesville, OK). After centrifugation at 10,000 rpm for 2 minutes, the supernatant was harvested and used for the measurement of flagellin, total IgA, and flic-specific IgA.

#### ELISA

96-well ELISA plates (Santa Cruz Biotechnology, Inc., Santa Cruz, CA) were coated with either purified goat-anti-mouse IgA at 1 ⌈l/well (clone: A90-103A, Bethyl Laboratories, Inc.) for total IgA detection, or 0.5 ⌈g/ml flagellin from Salmonella Typhimurium strain 14028 (Enzo Life Sciences, Inc., Farmingdale, NY) for the measurement of Flic-specific IgA. Plates were blocked with PBS containing 10% bovine serum albumin (BSA, Sigma, St. Louis, MO). Stool supernatants were prepared as mentioned above. The supernatants were either not diluted (for detection of flic-specific IgA) or diluted 1,500 fold with PBS (for detection of total IgA). 100 μl of resulting supernatant was loaded into each well. After incubation for 1 hour at 37°C (for flic-specific IgA) or room temperature (for total IgA), horseradish peroxidase-conjugated goat-anti-mouse IgA (Sigma, St. Louis, MO) was added to detect IgA. For the measurement of peroxidase activity, TMS peroxidase substrate solution (Sigma, St. Louis, MO) was used. After adding stopping solution, absorbance was measured at 450 nm with a Synergy H1 plate reader within 2 hours. The levels of anti-flic IgA were equivalents of a purified anti-mouse flic IgG binding.

#### Bioactive flagellin quantification

Flagellin was quantified by using HEK-Blue-mTLR5 cell line according to the manufacturer’s instructions (Invivogen, San Diego, CA). Briefly, fecal samples were prepared as mentioned above at the concentration of 0.1 mg/μl and incubated at 94°C for 10 minutes. After diluting 2,000 fold, 20 μl of sample was cocultured with HEK293-mTLR5 reporter cell line for 18 hours. The next day, cell culture supernatant was applied to HEK-Blue Detection medium (Invivogen, San Diego, CA). Alkaline phosphatase activity was measured at 630 nm on a Synergy H1 microplate reader.

#### Tissue harvesting and cell preparation

Bone marrow, spleen, and intestines were harvested. Single-cell suspensions were prepared (Felio et al., 2009; Weigmann et al., 2007). For bone marrows, cells were flushed out from tibia and femur. Single-cell suspension was harvested for flow staining. For spleen, tissues were mechanically dissociated into single-cell suspensions. For intestines, tissues were flushed with cold PBS, opened longitudinally, washed and cut into 0.5 cm pieces. Tissue fragments were treated at 37°C for 20 minutes with PBS containing 1 mM dithiothreitol and 20 mM EDTA to remove epithelial cells. The remaining tissues were then minced and dissociated at 37°C for 20 minutes with shaking at 100 rpm in PBS containing 0.5 mg/ml collagenase, 0.5 mg/ml DNase I (Roche Diagnostics) and 2% FBS. The digested suspensions were harvested, pelleted, and subjected to discontinuous 40% / 80% Percoll gradient separation. The resulting cells were collected from Percoll interface.

#### Flow cytometric staining

Single-cell suspensions were isolated as described above. 1-2 × 10^6^ cells were incubated with Fc blocking antibody for 15 minutes at 4°C, followed by incubation with anti-mouse mono-antibodies (eBioscience) for 30 minutes at 4°C: CD11b (ICRF44), Ly6G(Gr1, 1A8-Ly69), CD3(145-2C11), CD19 (1D3), mouse IgG2a, and TLR5(19D759.2). For IgA intracellular staining, LIVE/DEAD™ Fixable Violet Dead Cell Stain Kit (L34955, ThermoFisher, US) was used to distinguish live cells from dead cells according to the manufacturer's instructions. After surface stained with CD19, cells were fixed in 4% paraformaldehyde, permeabilized with 0.1% saponin and stained with anti-mouse IgA (mA-6E1). The data of flow cytometric staining were collected on a LSRII (BD) and analyzed using FlowJo (Tree Star).

#### Generation of bone marrow chimeras (BMC)

Five ten-week-old WT or TLR5^-/-^ male mice, derived from multiple breeders, were housed together after weaning and served as recipients. Recipients were sublethally irradiated with a total dose of 1,000 rad. 3-6 hours after irradiation, a total of 10 million bone marrow cells, which had been isolated from the tibias and femurs of 12-week old male donor WT and TLR5^-/-^ mice, were injected into the recipient mice retro-orbitally. Four chimeric groups were generated: WT > WT, WT > TLR5^-/-^, TLR5^-/-^ > WT, and TLR5^-/-^ > TLR5^-/-^ (donor>recipient), and placed on cherry flavored sulfamethoxazole and trimethoprim in drinking water (final concentration: 0.5 mg/ml and 0.1 mg/ml) for one week after irradiation. Body weights were monitored weekly. Fecal samples were collected twice a week. This experiment was performed twice (Fig 4 and Supplemental figure 1). Date from littermates and non-littermates (for both donor and recipients) were plotted separately or represented by different symbols.

#### Genotyping TLR5 in BMC recipients

Genotyping blood cells was performed as according to the instruction from the website of Jackson lab (https://www.jax.org/jax-mice-and-services/customer-support/technical-support/genotyping-resources/dna-isolation-protocols). Briefly, blood was collected from the murine tail vein. After lysis, nuclei were pelleted, suspended in 100 μl PBND (50 mM KCl, 10 mM Tris-HCl pH 8.3, 0.1mg/ml gelatin, 0.45% Nonidet P40, 0.45% Tween 20) with 60 μg/ml proteinase K, and incubated at 55°C for 1 hour. Proteinase K was heat-inactivated, and 1-2 μl DNA solution was used for PCR-genotyping. PCR cycle conditions were as follows: 94°C initial denaturation for 3 minutes; 35 cycles of 94°C for 30 seconds, 64°C annealing for 1 minute, 72°C extension for 1 minute; final single extension at 72°C for 2 minutes. The protocol of PCR using high-fidelity HotStart Taq polymerase (Invitrogen) were as follows (12 μl total): 2 μl of DNA, 0.15 μl of each primer (10 μM), 1.2 μl of HF buffer(10x), 0.6 μl of MgSO4, 1.66 μl of Betaine (5M), 0.15 μl of 10 mM dNTP, 0.08 μl HotStart Taq polymerase and 5.86 μl of H2O. The primers used in genotyping were as following: Common primer: 5’-TGAACAAACACTGCCTGCGTG-3’; WT reverse: 5’-AACACCACATCACAGCCTGAGG-3’; Mutant: 5’-GTGGGATTAGATAAATGCCTGCTC (Zhang et al., 2016).

#### 16S rRNA gene sequencing

Fecal samples were harvested from recipient mice of bone marrow chimera assays at various time-points and homogenized by a Mini-Beadbeater-24 on high for 2 minutes in the presence of beads. Genomic DNA was extracted from each sample by using the PowerWater DNA isolation kit as according to the manufacturer’s instruction (MoBio Laboratories Ltd, Carlsbad, CA). 16S rRNA genes were amplified with 515F and 806R primers for the V4 hypervariable region of the 16S rRNA gene (Supplemental Table 2). Primers included a unique 12-base barcode to tag PCR products from different samples. An insert of ‘CC’ was used as a linker between the barcode and the gene primer sequence. PCR using Easy-A high-fidelity enzyme was performed as follows: 95°C initial denaturation for 2 minutes; 25 cycles of 95°C for 30 seconds, 57°C annealing for 45 seconds, 72°C extension for 60 seconds; final single extension at 72°C for 7 minutes. PCR products were purified using Ampure magnetic purification beads (Agencourt, Danvers, MA) and quantified using Quant-iT PicoGreen dsDNA kit (Invitrogen, Carlsbad, CA). Samples were pooled in equimolar ratios and sequenced using the Illumina Miseq 2000 2 × 250 bp platform at the Genomic Facility in Biotechnology Resource Center at Cornell University.

#### Analysis of 16S rRNA gene sequences

Quantitative Insights into Microbial Ecology (QIIME) 1.9.1 was used to process and analyze the 16S rRNA gene sequence data (Caporaso et al., 2010). After stitching (fastq-join, ea-utils 1.1.2-537) and demultiplexing with default quality settings, sequences were then clustered to operational taxonomic units (OTUs) using open reference OTU picking with default settings, followed by the taxonomic classification using the August 2013 Greengenes reference database. Alpha diversity box-whisker plots and non-parametric Monte-Carlo permutation significance tests were generated from 10x rarefied data at 10,000 sequences per sample. Linear mixed model tests for differential OTU abundance was done using log-transformed OTU count data (even sampled at 18442 sequences/sample) and the R package LME4. The model was lme4::lmer(OTU_count~Treatment + Plate + (1|Assay) + Date + (1|ID)) where treatment is the mouse donor/recipient category, Plate is the plate identity from sample processing, Assay is the replicate assay 1 or 2, and ID is mouse identity (assay and mouse id are treated as non-independent, random effects). Pairwise significance testing was done using the lsmeans R package, with: lsmeans::lsmeans(df.model, pairwise~Treatment, mode = “kenward-roger”).

#### BAFF measurement

Contents were removed from the large intestine by flushing with cold PBS. The intestines were opened longitudinally and washed extensively; tissues were homogenized in PBS containing Halt protease inhibitor (PIERCE) for 20 seconds twice at 4°C. Samples were centrifuged at 4°C for 15 minutes at 14,000 rpm. The supernatants were used for BAFF measurement with mouse BAFF/BLyS/TNFSF13B immunoassay kit (R&D Systems, Quantikine) according to the manufacturer’s instruction. Briefly, samples were loaded into the mouse BAFF/Blys pre-coated wells. Enzyme-linked antibody specific for mouse BAFF/BLys were used to detect BAFF. The substrate solution was added to measure enzyme activity. After adding the stop solution, absorbance was measured by OD value at the wavelength of 450 nm, with the correction wavelength at 540 nm.

#### B cell transfer

Splenocytes were isolated from 12-week old WT and TLR5^-/-^ male littermates by mechanical dissociation. Splenic B cells were enriched by CD19-positive selection (Miltenyi Biotech). >95% of the enriched populations were CD19^+^ cells. 1 × 10^7^ purified cells were transferred into seven-week-old male μMT mice retro-orbitally. Recipients were single-housed. Feces were collected twice per week. Body weights were monitored two times per week. Total IgA and anti-flic specific IgA in fecal samples was measured by ELISA. Flagellin level was measured by HEK Blue-mTLR5 reporter cell line.

#### Microbiota transplantation

Fresh cecal contents from 12-week old WT and TLR5^-/-^ male littermates were suspended in anaerobic PBS by vortexing. After sitting at RT for 10 minutes, the supernatant was orally administered to 3-4-week-old male GF Swiss Webster animals once daily for two consecutive days. Transplanted mice were single maintained in sterile cages for three weeks. Body weight was monitored. The percentages of IgA-secreting B cells, flagellin-specific IgA level, and total IgA level were examined at the end of week three.

#### Ig Repertoire sequencing

Splenocytes and mononuclear cells from the lamina propria of the large intestine were isolated from TLR5^-/-^ mice and their WT littermates. Total RNA was isolated from cells using Trizol Reagent (Invitrogen) according to the manufacturer’s instructions. After phenol-chloroform extraction, RNAs were used to generate library following the modified protocol for a High-Throughput Illumina Strand-Specific RNA Sequencing Library preparation(Zhong et al., 2011). Briefly, 2 ⌈g of total RNA was used to enrich mRNA with magnetic oligo(dT) beads (Invitrogen). After fragmentation at 95°C for 5 minutes and elution from beads, cleaved RNA fragments were primed with the external reverse primer targeted IgA heavy-chain constant region to synthesize the first cDNA strand using reverse transcriptase SuperScript III (Invitrogen) with dNTP (Supplemental Table 1). Forward primers were designed based on alignments of IMGT reference sequences from B6 mice, and degenerate nucleotides were introduced to cover multiple alleles in one subclass (Supplemental Table 1). The synthesized cDNA was then split evenly and used as template in triplicate 25 ⌈l PCR reactions with the internal reverse primer positioned in a distinct IgA heavy-chain constant region, along with a mix of 18 forward primers (incorporated based on the abundance of their targets). The protocol of PCR using high-fidelity Phusion HotStart II polymerase were as follows: 2 μl of cDNA, 0.5 μl of reverse primers (10 μM), 0.5 μl of a mixed 18 forward primers, 5 μl of Phusion HF buffer(5x), 0.25 μl of 10 mM dNTP, 0.5 μl Phusion II and 7.75 μl of H2O. Cycle conditions were as follows: 98°C initial denaturation for 2 minutes; 15 cycles of 98°C for 20 seconds, 60°C annealing for 20 seconds, 72°C extension for 20 seconds; final single extension at 72°C for 2 minutes. Triplicate PCR reactions were then combined, blunt-repaired (Enzymatics), and added ‘A’ tailing at the ends (Klenow 3’-5’ exo-, Enzymatics). The barcoded adaptors were ligated at the 3’ end of ‘A’-tailed-dsDNA. PCR indexed primers annealing in the adaptor sequence were then used for the final library amplifications at 98°C initial denaturation for 30 seconds; 5 cycles of 98°C for 20 seconds, 65°C annealing for 30 seconds, 72°C extension for 30 seconds; final single extension at 72°C for 2 minutes. The libraries were pooled together (10 ng/library), purified with 75% ethanol, concentrated with XP beads (Beckman Coulter) and sent to the Genomics Facility at Biotechnology Resource Center at Cornell University for paired-end sequencing (2 x 300 bp) performed with Miseq Illumina platform.

#### Ig sequence analysis

Sequences were initially processed to create stitched (SeqPrep 1.1) reads that were demultiplexed and quality filtered with QIIME 1.9.1. A custom script was used to find and trim out the primer sequences (https://gist.github.com/walterst/2fce207ff38ad04c0bcbb2e8531ac230) and discard reads where either primer was not found. The prefix-suffix OTU picking method (--prefix_length 1000) of QIIME had applied to cluster identical sequences. Based on the grouped sequences, the biom table of counts of Ig sequences by samples for each repertoire was created and used to calculate mean observed OTUs and Shannon alpha diversity values at 200000 sequences per sample, randomly subsampled 10 times. Statistic significance was evaluated via a nonparametric permutation test with 999x repeats via QIIME’s compare_alpha_diversity.py script‥ For beta diversity, IgA read data were evenly sampled at 200000 sequences per sample. Bray-Curtis dissimilarity measures were calculated on this rarefied OTU table and used for generating PCoA plots and calculating Adonis results (QIIME’s compare_categories.py script). Kruskal-Wallis tests (QIIME’s group_significance) were performed on the same even-sampled OTU table for differential abundance by genotype after splitting the OTU table by organ (spleen and large intestine). Rank and frequency sorted data for high and low frequency IgA population determination was accomplished by first sorted the even-sampled OTU table data with a custom script (https://gist.github.com/walterst/ecc2232c07d2011d7fdd2c06fb0071a8). The output from this custom script was imported into Microsoft Excel, and this equation was used to classify data as high or low frequency: Xc = e^((log N + log a)/−b). N is the number of different CDR3 sequences, and a and b are the intercept and slope (respectively) of the log-log plot. All ranks belonging to the power law component (the low frequency clones) must have a smaller rank than Xc.

#### Statistical analysis

Data were represented as mean ± S.D. Statistical significance was evaluated using a two-tailed ANOVA test with a 95% confidence interval. The analysis was performed with a Prism 7 program (GraphPad Software).

Sequence data were deposited in the European Nucleotide Archive under the studies PRJEB14726 (IgA receptor sequences) and PRJEB14730 (16S ribosomal small subunit gene amplicons).

## Acknowledgements

The authors thank Jessica Sutter and Noah Clark for technical assistance in the preparation of 16S rRNA gene libraries and the maintenance of germ-free mice. The authors also thank the Genomics Facility at the Biotechnology Resource Center and Flow cytometry core facility at Cornell University. This work was supported by grant by a NIH Director’s New Innovator Award (DP2 OD007444) and the Max Planck Society.

## Supplemental Figures

**Supplemental Fig. 1:**
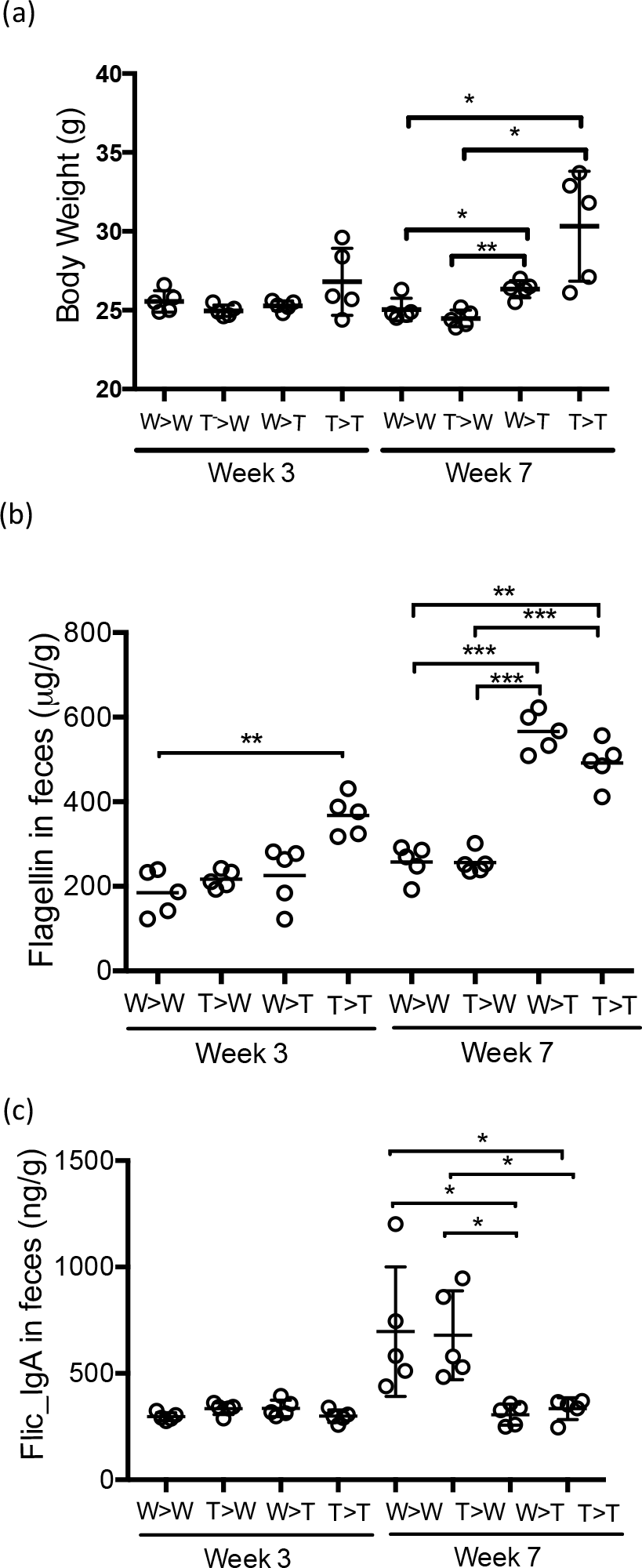
TLR5 on non-hematopoietic cells maintains healthy metabolism and gut homeostasis. Replicate sample results (see Figure 4). Age-matched WT and TLR5^-/-^ littermates were used in this assay for both donor and recipients with n = 5 recipients/group. (a) Body weight. (b) Fecal flagellin level. (c) Flic-specific IgA concentration. **: p<0.01; ***: p<0.0001; n.s.: not significant.

**Supplemental Fig. 2:**
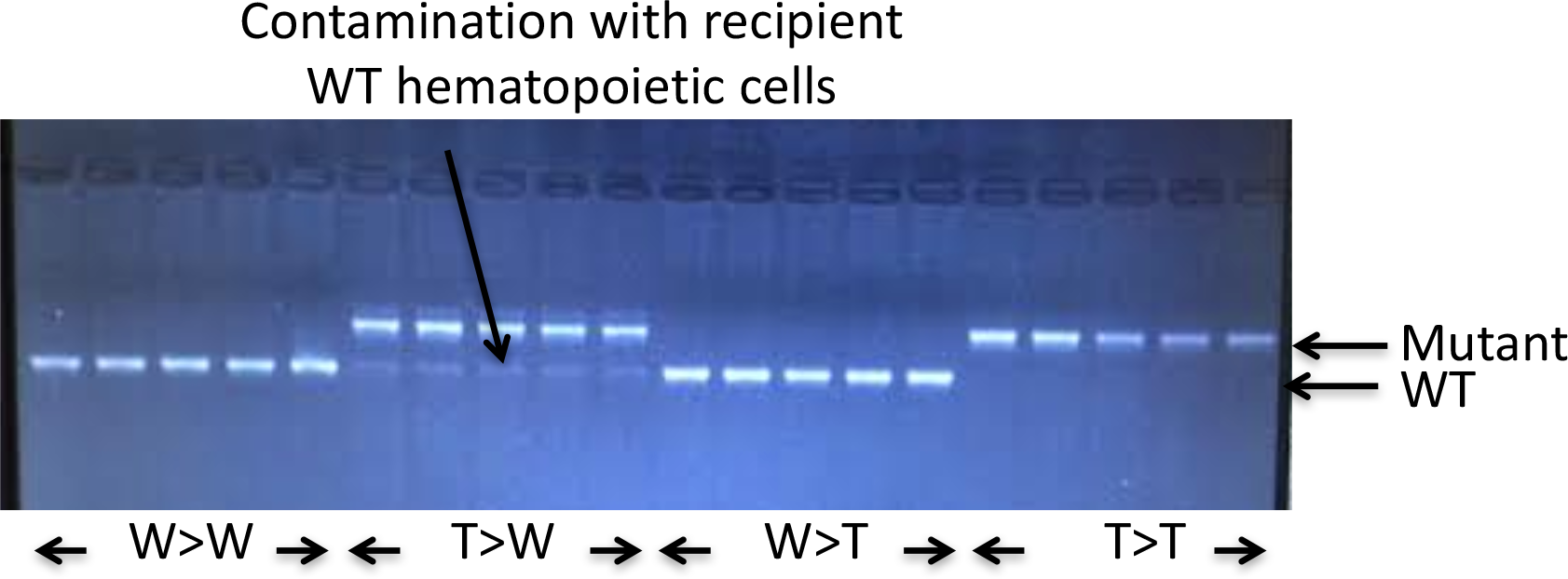
Contamination of recipient hematopoietically-differentiated cells in the chimera group of TLR5^-/-^ to WT. Blood was harvested from all recipient mice at week four after bone marrow transfer. Genomic DNAs were isolated from blood lymphocytes. WT and mutant TLR5 genes were PCR amplified with gene-specific primers and 2% agarose gel electrophoresis was used to separate WT and mutant TLR5 product. Data were representative of two independent assays.

**Supplemental Fig 3:**
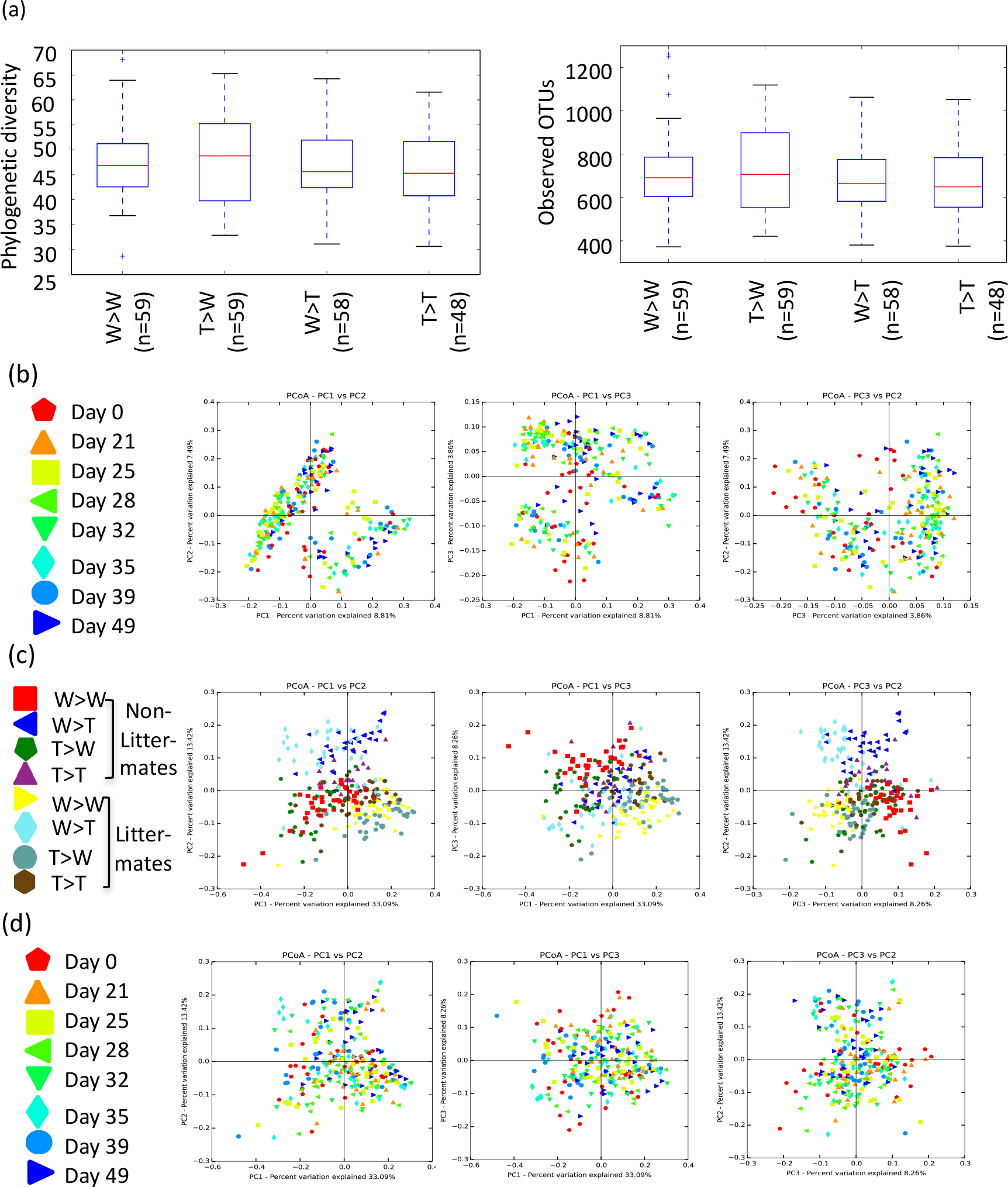
Microbiome alpha diversity and beta diversity for chimeric WT and TLR5^-/-^ mice. (a) Faith’s Phylogenetic diversity and Observed OTUs alpha diversity measures for each donor and recipient category (no significant differences). (b) Unweighted UniFrac based clustering of samples, colorized by time point. (c) Weighted UniFrac clustering of samples, colorized by donor/recipient genotype and littermate status. (d) Weighted UniFrac clustering of samples, colorized by time gradient.

**Supplemental Fig. 4:**
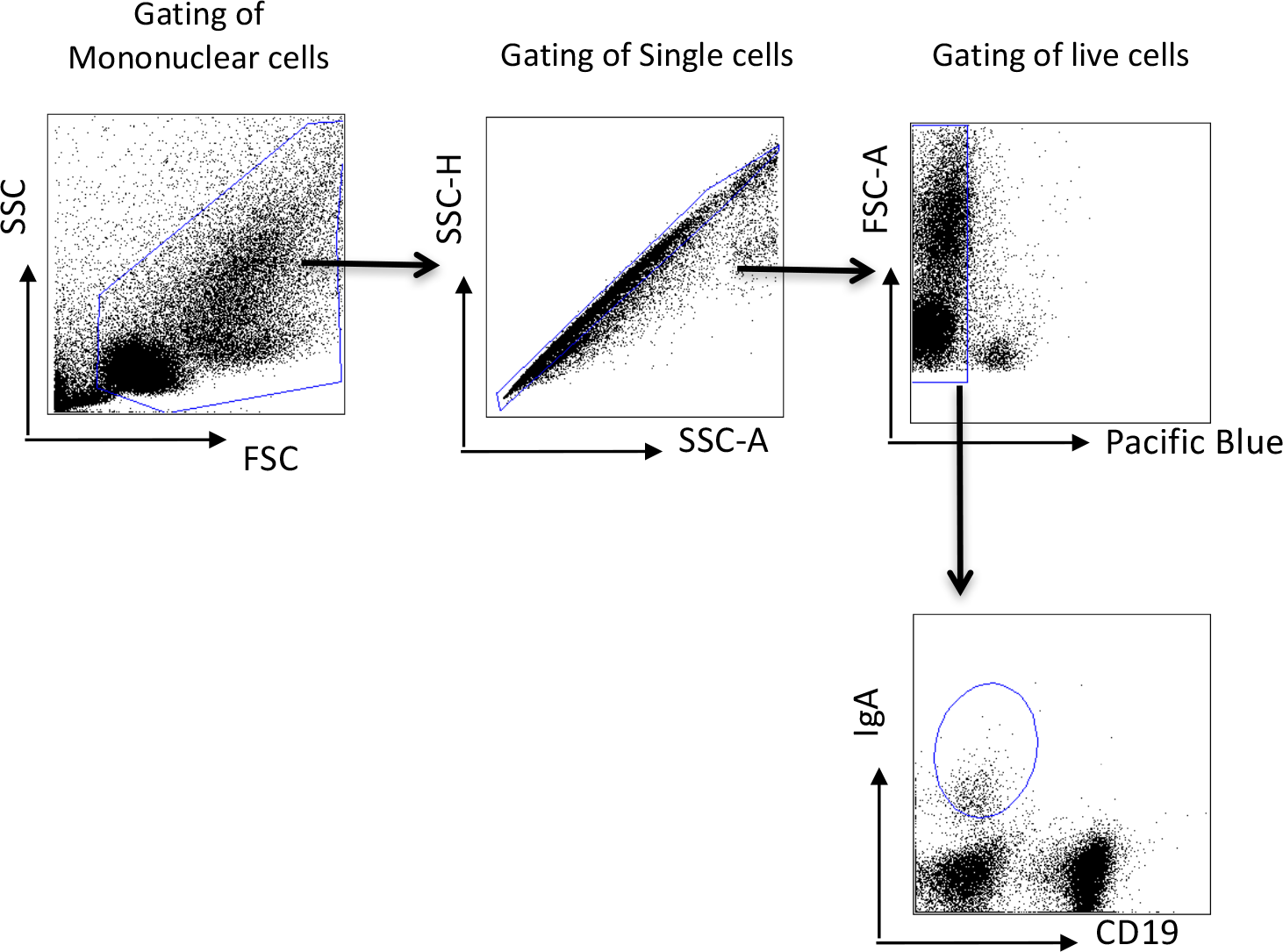
A gating scheme for IgA-secreting cells. The gating strategy shown illustrates the elimination of doublets and dead cells.

**Supplemental Table 1:** Linear mixed model results for differential abundance of 16S based OTUs for chimeric mice. Mice are combinations of WT and TLR5−/− bone marrow donors and recipients, for a total of four categories. OTU data were log transformed with the four categories, sample plate, and date as factors, while mouse ID and assay were treated as a random effects. Bonferroni correction used for p-values for number of OTUs tested. q-q plots and residual histograms deviating from normal distributions were filtered from results.

| Genotype 1 | Genotype 2 | q-value | Mean sequence count, genotype 1 | Mean sequence count, genotype 2 | Taxonomy (Greengenes 13_8) | OTUId |
| --- | --- | --- | --- | --- | --- | --- |
| WT->WT | WT->TLR5 | 5.01E-06 | 28.86292752 | 2.646591824 | k__Bacteria;p__Firmicutes;c__Clostridia;o__Clostridiales;f__Clostridiaceae;g__S__ | 555945 |
| WT->WT | TLR5->TLR5 | 0.034008104 | 28.86292752 | 6.780329204 | k__Bacteria;p__Firmicutes;c__Clostridia;o__Clostridiales;f__Clostridiaceae;g__S__ | 555945 |
| TLR5->WT | WT->TLR5 | 1.15E-07 | 44.45018423 | 2.646591824 | k__Bacteria;p__Firmicutes;c__Clostridia;o__Clostridiales;f__Clostridiaceae;g__S__ | 555945 |
| TLR5->WT | TLR5->TLR5 | 0.000614416 | 44.45018423 | 6.780329204 | k__Bacteria;p__Firmicutes;c__Clostridia;o__Clostridiales;f__Clostridiaceae;g__S__ | 555945 |
| WT->WT | WT->TLR5 | 0.002547435 | 59.18383431 | 8.564006999 | k__Bacteria;p__Firmicutes;c__Clostridia;o__Clostridiales;f__Clostridiaceae;g__S__ | 780650 |
| TLR5->WT | WT->TLR5 | 0.000954081 | 66.71067778 | 8.564006999 | k__Bacteria;p__Firmicutes;c__Clostridia;o__Clostridiales;f__Clostridiaceae;g__S__ | 780650 |
| TLR5->TLR5 | WT->TLR5 | 0.044086484 | 42.73483984 | 8.564006999 | k__Bacteria;p__Firmicutes;c__Clostridia;o__Clostridiales;f__Clostridiaceae;g__S__ | 780650 |
| TLR5->WT | WT->WT | 4.98E-05 | 70.54521416 | 1.785922104 | k__Bacteria;p__Firmicutes;c__Clostridia;o__Clostridiales;f__Ruminococcaceae;g__S__ | 1110191 |
| TLR5->TLR5 | WT->WT | 0.02817996 | 22.23032304 | 1.785922104 | k__Bacteria;p__Firmicutes;c__Clostridia;o__Clostridiales;f__Ruminococcaceae;g__S__ | 1110191 |
| WT->WT | WT->TLR5 | 1.47E-05 | 351.8499982 | 19.00099327 | k__Bacteria;p__Firmicutes;c__Clostridia;o__Clostridiales;f__g__S__ | 268220 |
| TLR5->WT | WT->TLR5 | 1.71E-05 | 344.0541437 | 19.00099327 | k__Bacteria;p__Firmicutes;c__Clostridia;o__Clostridiales;f__g__S__ | 268220 |
| TLR5->TLR5 | WT->TLR5 | 0.000313791 | 234.7942829 | 19.00099327 | k__Bacteria;p__Firmicutes;c__Clostridia;o__Clostridiales;f__g__S__ | 268220 |
| WT->WT | WT->TLR5 | 2.67E-06 | 59.36975098 | 9.977324454 | k__Bacteria;p__Bacteroidetes;c__Bacteroidia;o__Bacteroidales;f__S24-7;g__S__ | 276509 |
| TLR5->WT | WT->TLR5 | 1.16E-06 | 63.25691561 | 9.977324454 | k__Bacteria;p__Bacteroidetes;c__Bacteroidia;o__Bacteroidales;f__S24-7;g__S__ | 276509 |
| TLR5->TLR5 | WT->TLR5 | 1.64E-09 | 113.5880875 | 9.977324454 | k__Bacteria;p__Bacteroidetes;c__Bacteroidia;o__Bacteroidales;f__S24-7;g__S__ | 276509 |
| WT->WT | WT->TLR5 | 8.42E-07 | 13.76459882 | 1.06150495 | k__Bacteria;p__Bacteroidetes;c__Bacteroidia;o__Bacteroidales;f__S24-7;g__S__ | 198964 |
| WT->WT | TLR5->TLR5 | 6.50E-07 | 13.76459882 | 1.03492587 | k__Bacteria;p__Bacteroidetes;c__Bacteroidia;o__Bacteroidales;f__S24-7;g__S__ | 198964 |
| TLR5->WT | WT->TLR5 | 2.72E-11 | 47.28462327 | 1.06150495 | k__Bacteria;p__Bacteroidetes;c__Bacteroidia;o__Bacteroidales;f__S24-7;g__S__ | 198964 |
| TLR5->WT | TLR5->TLR5 | 6.69E-12 | 47.28462327 | 1.03492587 | k__Bacteria;p__Bacteroidetes;c__Bacteroidia;o__Bacteroidales;f__S24-7;g__S__ | 198964 |
| TLR5->TLR5 | TLR5->WT | 0.005465373 | 78.98547435 | 12.25303185 | k__Bacteria;p__Bacteroidetes;c__Bacteroidia;o__Bacteroidales;f__S24-7;g__S__ | 258849 |
| WT->WT | TLR5->TLR5 | 0.007514674 | 42.21278165 | 2.900413431 | k__Bacteria;p__Bacteroidetes;c__Bacteroidia;o__Bacteroidales;f__S24-7;g__S__ | 270984 |
| WT->TLR5 | TLR5->TLR5 | 0.000414255 | 70.70035574 | 2.900413431 | k__Bacteria;p__Bacteroidetes;c__Bacteroidia;o__Bacteroidales;f__S24-7;g__S__ | 270984 |
| TLR5->WT | TLR5->TLR5 | 0.001439779 | 56.69009906 | 2.900413431 | k__Bacteria;p__Bacteroidetes;c__Bacteroidia;o__Bacteroidales;f__S24-7;g__S__ | 270984 |
| TLR5->TLR5 | WT->WT | 0.029136291 | 24.26763983 | 9.82318352 | k__Bacteria;p__Tenericutes;c__Mollicutes;o__RF39;f__g__S__ | 2286580 |
| WT->TLR5 | TLR5->WT | 0.008495051 | 17.05658067 | 6.51335281 | k__Bacteria;p__Tenericutes;c__Mollicutes;o__RF39;f__g__S__ | 2286580 |
| TLR5->TLR5 | TLR5->WT | 5.82E-05 | 24.26763983 | 6.51335281 | k__Bacteria;p__Tenericutes;c__Mollicutes;o__RF39;f__g__S__ | 2286580 |
| A total of 10 OTUs had significant results, for many of these, more than one comparison between genotypes is significant, for a total of 26 significant differences |  |  |  |  |  |  |
| Data were log transformed to better fit assumptions underlying models. Results with q-q plots or histograms that reflected non-normal data were filtered by hand |  |  |  |  |  |  |
| This was the model generated in R: |  |  |  |  |  |  |
| df.model <- lme4::lmer(LogXform~Treatment + Plate + (1\|Assay) + Date + (1\|ID)) |  |  |  |  |  |  |
| There was a strong assay effect when visualizing clusters in beta diversity, so this is set as a dependent effect, and the mouse itself that is repeatedly sampled (ID) is also non-independent. |  |  |  |  |  |  |
| Followed by a pairwise test: |  |  |  |  |  |  |
| pairwise_results = lsmeans::lsmeans(df.model, pairwise~Treatment, mode = "kenward-roger") |  |  |  |  |  |  |
| There are four sample categories considered, and these are based upon the two donor types (WT or TLR5) and recipients (WT or TLR5). |  |  |  |  |  |  |
| These are annotated with Donor->Recipient, e.g. WT->TLR5 or WT->WT |  |  |  |  |  |  |

**Supplemental Table 2:**
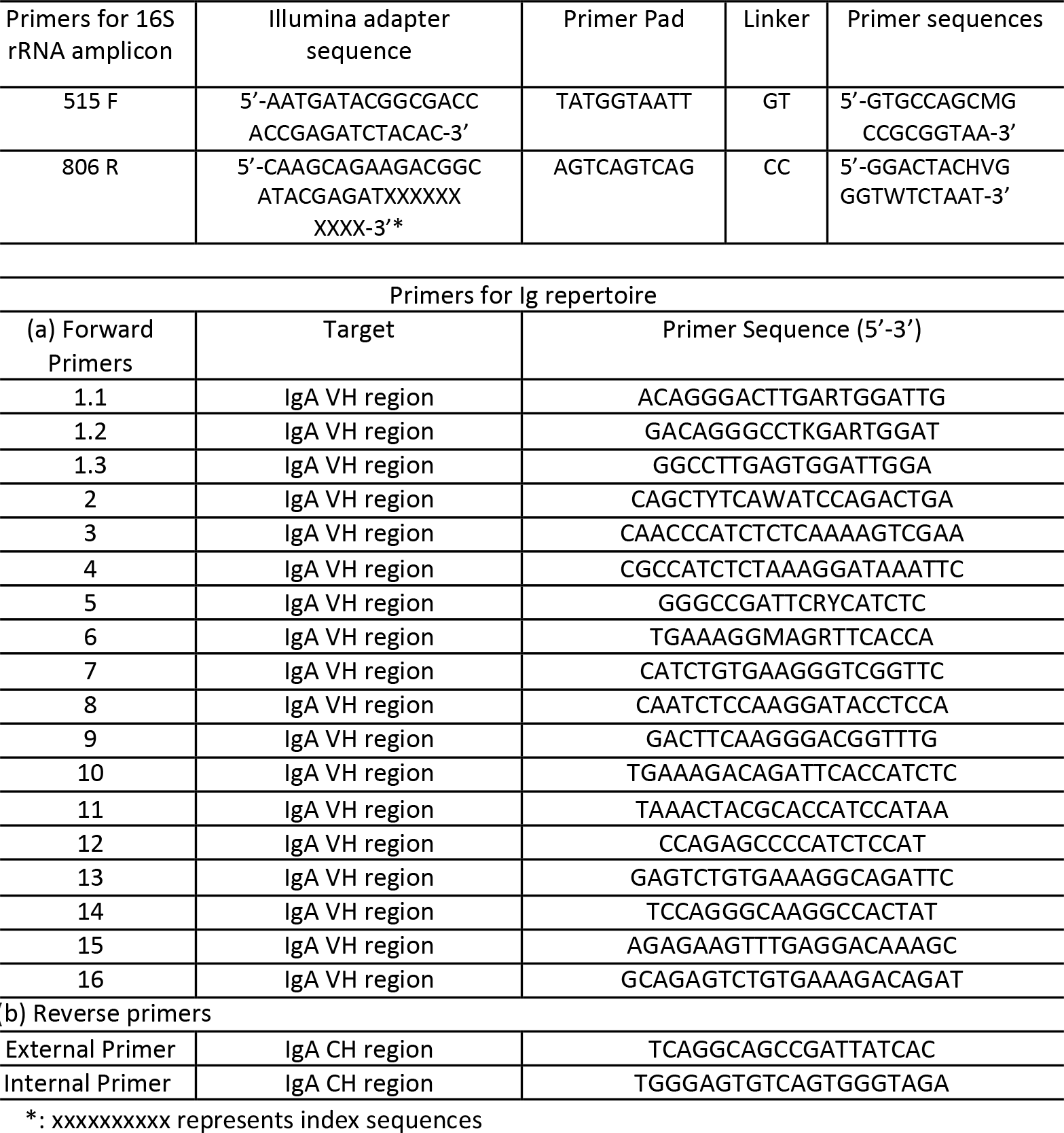
Primers used in sequencing 16s rRNA SSU gene and IgA repertoire

**Supplemental Table 3:** IgA CDR3 sequence reads per sample

| Organ | Number of sequence reads* |  |
| --- | --- | --- |
|  | WT | Tlr5<sup>-/-</sup> |
| Spleen | 459,000 ± 153,000 | 360,000 ± 113,000 |
| Lamina propria in the large intestine | 366,000 ± 117,000 | 380,000 ± 121,000 |
| * Data are represented as mean ± standard deviation |  |  |

**Supplemental Table 4:** Adonis analysis of CDR3 region of IgA heavy chain. R^2^ and p-values were calculated on each category using a Bray-Curtis dissimilarity matrix which was calculated from data evenly sampled to 200000 sequences per sample

| Category | R<sup>2</sup> | p-value |
| --- | --- | --- |
| Mouse | 0.82841 | 0.001 |
| Genotype | 0.18888 | 0.004 |
| Genotype X Organ | 0.23666 | 0.411 |
| Organ | 0.02482 | 0.961 |

